# A conserved node degree-based backbone and flexible hub organization of brain connectome underlying naturalistic movie watching

**DOI:** 10.64898/2026.03.09.710639

**Authors:** Xuehu Wei, Laura Rigolo, Colin P. Galvin, Einat Liebenthal, Yanmei Tie

## Abstract

How does the brain organize and transmit information during naturalistic conditions? Using large-scale movie-watching fMRI data spanning 14 diverse clips, validated in an independent dataset, we revealed a degree-based dual architecture of the functional connectome under naturalistic stimuli. During movie watching, the brain consistently expresses a conserved network backbone characterized by persistently high node degree in temporal and occipital sensory cortices and parietal association regions, whereas anterior higher-order cognitive regions show relatively lower node degree and substantially greater variability across different clips. We further found that this backbone links brain network organizational patterns to stimulus features of the clips, particularly audiovisual features capturing human presence and social communication. Within this backbone, subregions in superior temporal gyrus (STG), temporo-parieto-occipital junction (TPOJ), precuneus/posterior cingulate cortex, intraparietal sulcus (IPS), dorsal/ventral visual stream (DVS/VVS), and middle temporal visual and motion-sensitive areas (MT+) frequently emerge as connetome hubs, statistically mediating associations between multiple stimulus features and the integration of distinct large-scale functional systems. This study demonstrated a mechanistic framework in which movie watching relies on a conserved network backbone for stable large-scale integration, alongside stimulus-dependent rich-club hubs that flexibly mediate naturalistic information across distributed cortical systems.

---

Nervous systems operate as networks that transmit and process information ^1,2^, with communication and co-activation across brain regions forming the connectome that underlies cognitive processing ^3,4^. During real-life experiences, the brain coordinates activity across distributed regions to process complex information, supporting two core functions: transforming sensory inputs into representations of the external world ^5,6^, and integrating these inputs with internal states, such as intentions, motivations, and predictions, to construct coherent perceptual interpretations ^7^. Such integration is thought to rely on coordinated interactions within large-scale brain networks, yet how these interactions are organized under naturalistic conditions remains incompletely understood.

Movie watching has emerged as a widely adopted naturalistic paradigm for studying brain function ^8–10^. Movies are considered naturalistic stimuli because they integrate diverse visual elements, sounds, speech, music, social interactions, and narrative structure, evoking robust and reliable brain activity representative of natural cognition ^11,12^. Consistent with this view, movie stimuli elicit large-scale neural responses across occipital, temporal, and parietal cortices, inferior frontal and orbital prefrontal regions, as well as subcortical structures ^8,9,13^. These regions support hierarchical sensory processing information ^8,14^, language and narrative comprehension ^15^, social and emotional memory functions ^16,17^, and context integration and sustained attention ^18^, highlighting the brain’s capacity to coordinate multiple specialized systems during diversitial, real-world perception.

Despite the engagement of shared large-scale networks, neural responses to movies are not identical across stimuli. Variations in dialogue, narrative structure, and social content shape both the strength and spatial extent of cross-subject neural synchronization ^13,19,20^. Movies with rich audiovisual content and coherent narratives evoke widespread synchronization extending into higher-order association regions such as the inferior parietal lobe and posterior cingulate / precuneus ^8,21^, whereas stimuli lacking specific modalities or narrative coherence elicit responses in fewer regions ^22^. Together, these findings indicate that stimulus content strongly shapes neural responses. However, most prior work has emphasized response magnitude and regional engagement, leaving open whether diverse naturalistic stimuli are supported by a shared large-scale network framework, and which network mechanisms enable content-dependent information trasfer across distributed brain systems.

Addressing these questions requires moving beyond region-wise response strength to a network-level description of how distributed interactions are organized. Network neuroscience provides a framework for addressing this gap by characterizing brain function in terms of graph-theoretical properties such as node degree and hub regions ^23,24^. Among these properties, rich-club hubs, i.e., highly connected regions that form a central backbone for global information communication, are thought to support the integration of information across functional systems ^25,26^. While rich-club organization is well established in resting-state and task-based networks ^27^, recent work has begun to extend this graph-theoretical framework to naturalistic paradigms ^28,29^, probing whether hub-based backbones also support information integration during real-world perception.

In this study, we addressed these questions by analyzing movie-watching functional magnetic resonance imaging (fMRI) data from the Human Connectome Project (HCP), complemented by an independent in-house dataset. The combination of rigorous image quality, ample sample size, and stimulus heterogeneity provides rich variability and a solid empirical foundation for characterizing the general and specific topological features of brain network organization in response to different movie clips. We constructed stimulus-locked functional networks using inter-subject functional correlation (ISFC) ^30,31^ and applied data-driven graph-theoretical modeling ^32^ to characterize large-scale organizational regularities as well as clip-specific reconfigurations of network topology. Specifically, we first asked which large-scale network properties remain conserved across distinct movie clips during naturalistic viewing, and whether such stability reflects a common backbone architecture. We further identified the rich-club hubs and investigated whether the recruitment and connectivity patterns of these hubs vary systematically with stimulus content demands ^33–35^. Finally, we examined how multidimensional properties of movie stimuli relate to large-scale network organization, and whether rich-club hubs mediate the association between stimulus features and distributed cortical networks.

## Results

We used movie-watching fMRI data from 176 healthy young adults from HCP database (https://www.humanconnectome.org/study/hcp-young-adult) to investigate the characteristics of brain functional connectome in response to diverse short movie clips. fMRI data were acquired using a 7T scanner while participants underwent four movie-watching runs (Movies 1-4, Fig. 1A). Each run consisted of 3–4 short clips excerpted from either independent films (Movie 1 and Movie 3) or Hollywood movies (Movie 2 and Movie 4; see Methods and Supplementary Table 1 for details), and ended with the same validation clip. We conducted a standardized brain network analysis, building on ISFC in combination with complex network analysis (analysis details are provided in the Methods section and illustrated in Fig. 1B). ISFC was constructed using fMRI time-series data extracted from 374 regions-of-interest (ROIs), including 360 cortical regions defined by the HCP MMP 1.0 atlas ^36^ and and 14 subcortical regions defined by the Harvard-Oxford atlas ^37^. Fig. 2A shows the group-averaged ISFC matrix of each movie clip.

**Fig 1.**
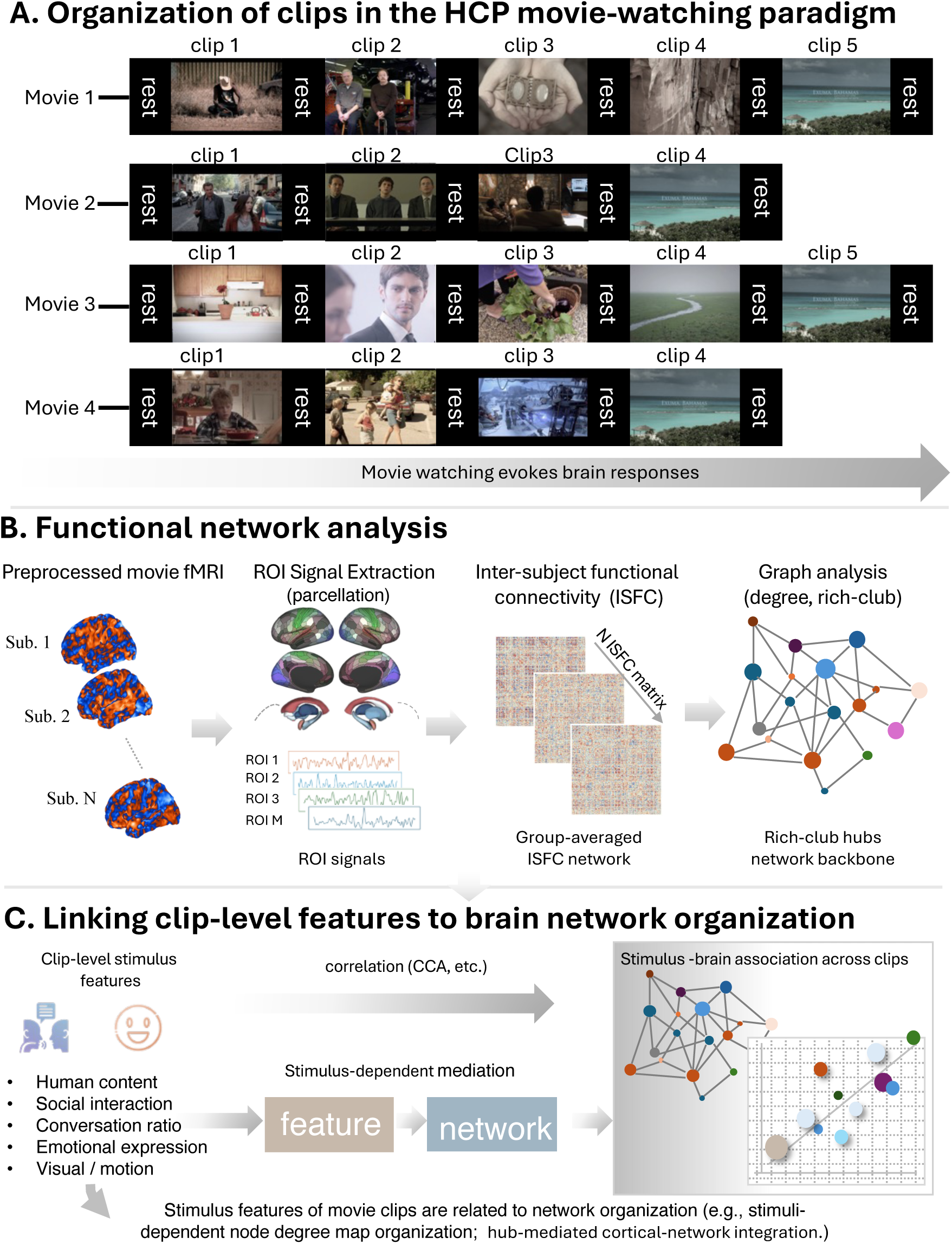
Movie-watching fMRI stimuli and data analysis workflow. (A) Organization of movie clips in the Human Connectome Project (HCP) movie-watching paradigm. Participants underwent four movie fMRI runs composed of multiple clips (about·1-4 minutes) presented in a fixed order and interleaved with brief rest periods (20 seconds). (B) Functional network analysis workflow. Preprocessed fMRI data were parcellated into cortical regions-of-interest (ROIs), and inter-subject functional connectivity (ISFC) was computed for each movie clip to capture stimulus-locked functional synchronization across individuals. Group-averaged ISFC networks were then analyzed using graph-theoretical measures, including node degree and rich-club organization, to identify integrative hub regions forming a core network backbone during movie watching. (C) Linking clip-level features to brain network organization. Clip-level visual semantic, auditory and social (e.g., face number, social interaction strength, and visual motion) were related to large-scale network measures using multivariate analyses. Feature–network mediation analyses further assessed how rich-club hub–to–network connectivity supports feature-specific processing within each cortical network.

**Fig 2.**
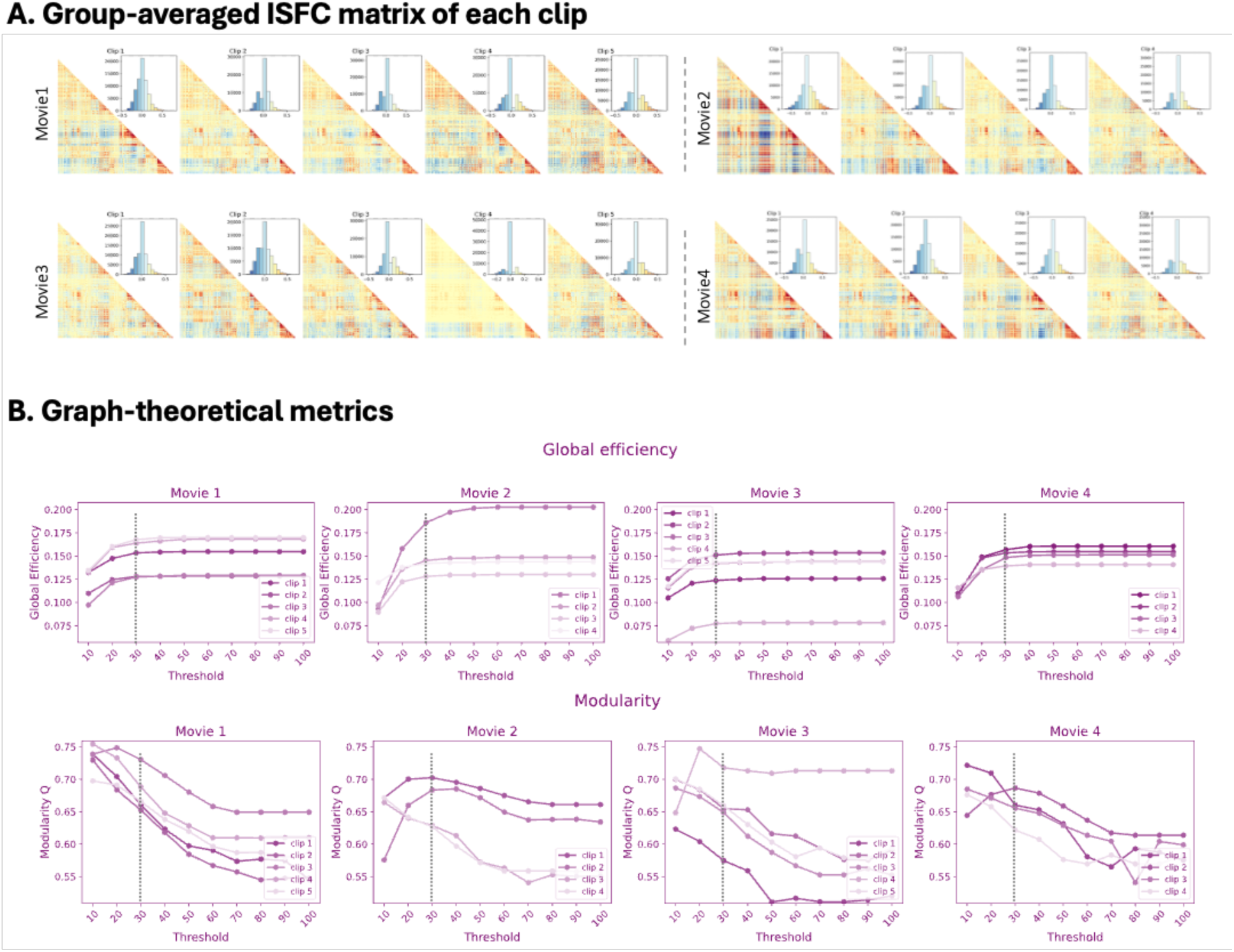
Group-averaged ISFC network properties. (A) Group-averaged ISFC matrix of each clip after removing false-positive connections through t-test verification. For each clip, the upper-triangular ISFC matrix is shown with the distribution of edge weights (inset). (B) Global graph-theoretical properties of clip-specific functional connectomes across a range of network density thresholds. Global efficiency (upper panel) and modularity (lower panel) are plotted for each clip within each movie run, demonstrating overall stability of global topology across threshold choices.

### Robust global network properties across density thresholds justify a common analysis regime

We began at the global level by evaluating key graph properties of the whole-brain functional connectome derived from group-averaged ISFC across a range of proportional density thresholds, constructed by retaining the strongest connections at each density level, focusing on global efficiency and modularity (Fig. 2). While all movie clips exhibited highly similar trends across density thresholds, the absolute values of these metrics varied across clips at each threshold (Fig. 2B; Supplementary Tables 2–3), consistent with differences in stimulus content. Global efficiency (upper panel of Fig. 2B), which indexes the capacity for large-scale functional integration during movie watching, showed clip-dependent variability, with maximum differences of 0.04, 0.07, 0.08, and 0.02 for movie runs 1–4, respectively. Modularity (lower panel of Fig. 2B), reflecting the extent to which the functional connectome is organized into segregated communities, also varied across clips, with differences of 0.10, 0.12, 0.20, and 0.08 observed for movie runs 1–4, respectively. The global network metrics of all movie clips (Fig. 2B) exhibited consistent trends with increasing thresholds above a 30% density threshold. Following the principle of economical small-world organization of graph theory ^38^, i.e., to support efficient integration at low cost while preserving modularity (Fig. 2B), we retained the top 30% of the strongest connections for all subsequent network analyses.

### Node degree-based connectome backbone is conserved across diverse movie clips

Having established a comparable network definition based on ISFC, we next examined whether the regional node degree patterns of the large-scale functional connectome remain conserved across clips. We quantified the node degree of each ROI (total 374 ROIs), i.e., the number of connections between an ROI and the rest of the brain that represents large-scale functional integration ^24^, for each of the 14 primary clips. The results indicated that the whole-brain node degree distribution patterns did not differ across clips (one-way repeated-measures ANOVA across clips, treating regions as samples; p > 0.05). Across all clips, node degree showed a highly consistent spatial distribution, with consistently elevated values in occipital cortex, dorsal and ventral visual streams (DVS and VVS), the temporo-parieto-occipital junction (TPOJ), and bilateral superior and middle temporal gyri (STG/MTG), whereas anterior and medial higher-order association regions showed comparatively lower node degree (Fig. 3A; Supplementary Fig. 1A). Notably, the node degree of STG/MTG increased in dialogue-rich clips and decreased in clips without speech (e.g., Movie 3 clips 1 and 4), yet consistently remained higher degree value than most higher-order regions. This pattern suggests that the stimulus-dependent modulation superimposed on a conserved backbone.

**Fig 3.**
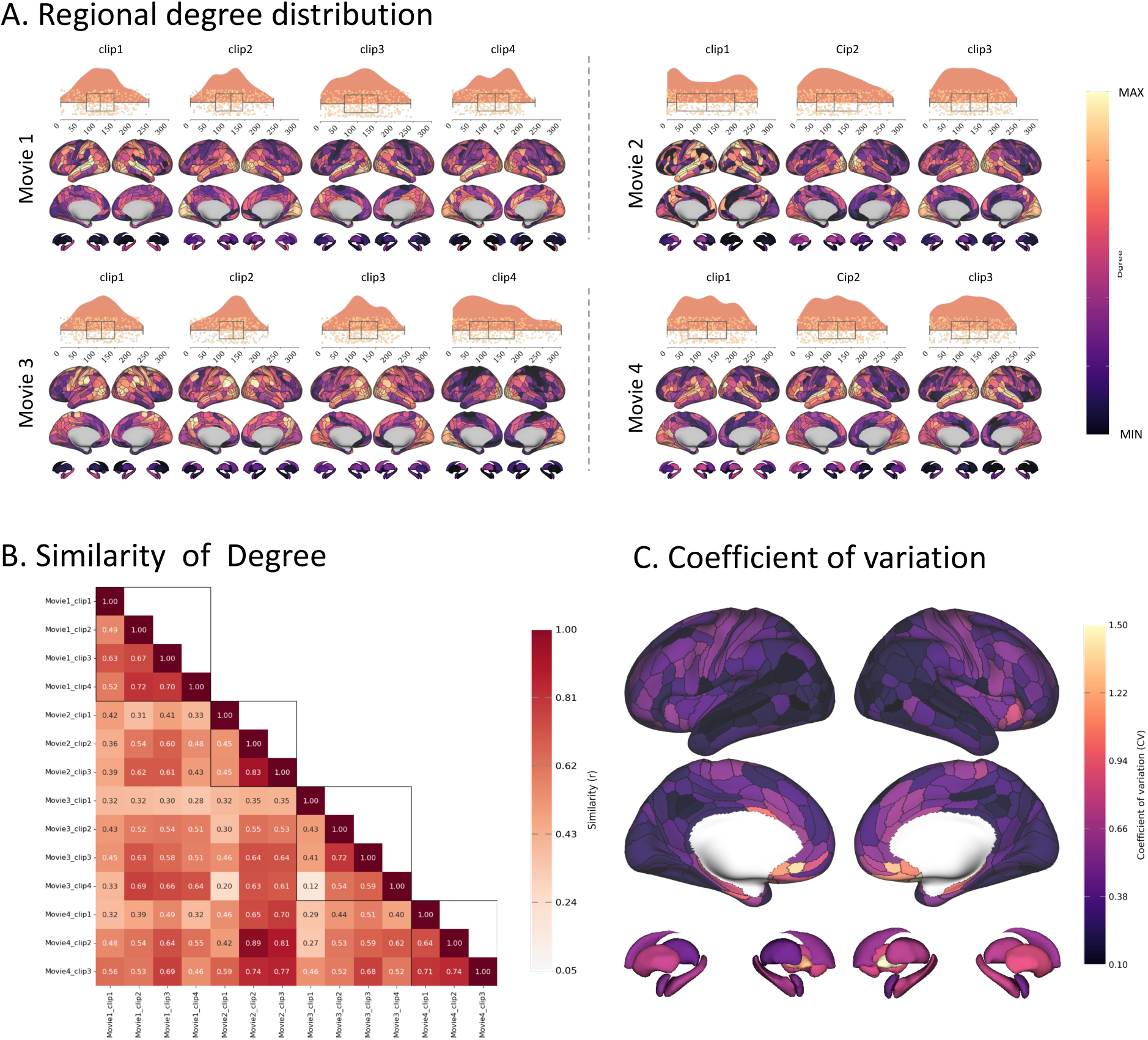
Stable node degree-based backbone across clips. (A) Spatial distribution of graph metric of node degree across 374 brain regions of each movie clip. For each clip, graph metrics of each region are shown on the brain maps with color indicating the magnitude of the metrics. Violin-Boxplots show the distribution of the metrics across all brain regions. (B) Similarity (Pearson’s correlation 𝑟) of node degree across movie clips. (C) Coefficient variation (CV) of node degree in each brain region across movie clips.

To quantify cross-clip stability, we computed pairwise Pearson correlations on the whole-brain node degree maps between each pair of the clips. Node degree distribution patterns were highly similar across many clip pairs (Fig. 3B), with strong within-run similarity (Movie 1: r = 0.49 - 0.72; Movie 2: r = 0.45 - 0.83; Movie 3: r = 0.12 - 0.72; and Movie 4: r = 0.64 - 0.74). Cross-run similarity was higher among Hollywood clips (Movies 2 and 4; mean r = 0.698) than that of independent clips (Movies 1 and 3; mean r = 0.493). The coefficient of variation (CV) of node degree across clips revealed that node degree variability differed substantially (ranging from 0.14 to 1.25) across brain regions (Fig. 3C; Supplementary Tables 4). As shown in Fig. 3C, variability was greater in orbitomedial frontal and parahippocampal regions, whereas occipital and posterior temporal regions exhibited comparatively lower variability across clips. This finding suggests that different brain regions expressed different levels of sensitivity to clip-specific content within a conserved node degree-based network backbone.

### Stimulus-dependent modulation of the node degree-based backbone aligns with latent movie feature dimensions

We next examined whether clip-to-clip variability in the organization of node degree maps was associated with systematic differences in stimulus features. To characterize moment-to-moment stimulus structure, we leveraged TR-resolved semantic feature annotations from the Human Connectome Project, which were generated using WordNet-based labeling of visual content. For visualization, we summarized the WordNet-based visual labels using word clouds (Fig. 4A; Supplementary Fig. 2A). In addition, we extracted speech and dialogue markers from the audio track using pyannote, and quantified the number of people appearing in each segment using YOLO-based detection. Together with eye-gaze annotations from the visual features, these measures were used to derive a time-resolved index of social interaction intensity (Fig. 4A; Supplementary Fig. 2B). These clip-level features provided a multidimensional description of clip content spanning both non-social visual content (e.g., objects, locations, and motion-related categories) and human-related cues (e.g., faces, speech, and social strength) (Fig. 4B; Supplementary Fig. 3). These features enabled systematic analyses linking naturalistic stimuli to graph-theoretical metrics (specifically node degree) within clip-specific functional connectomes, i.e., stimulus-brain association.

**Fig 4.**
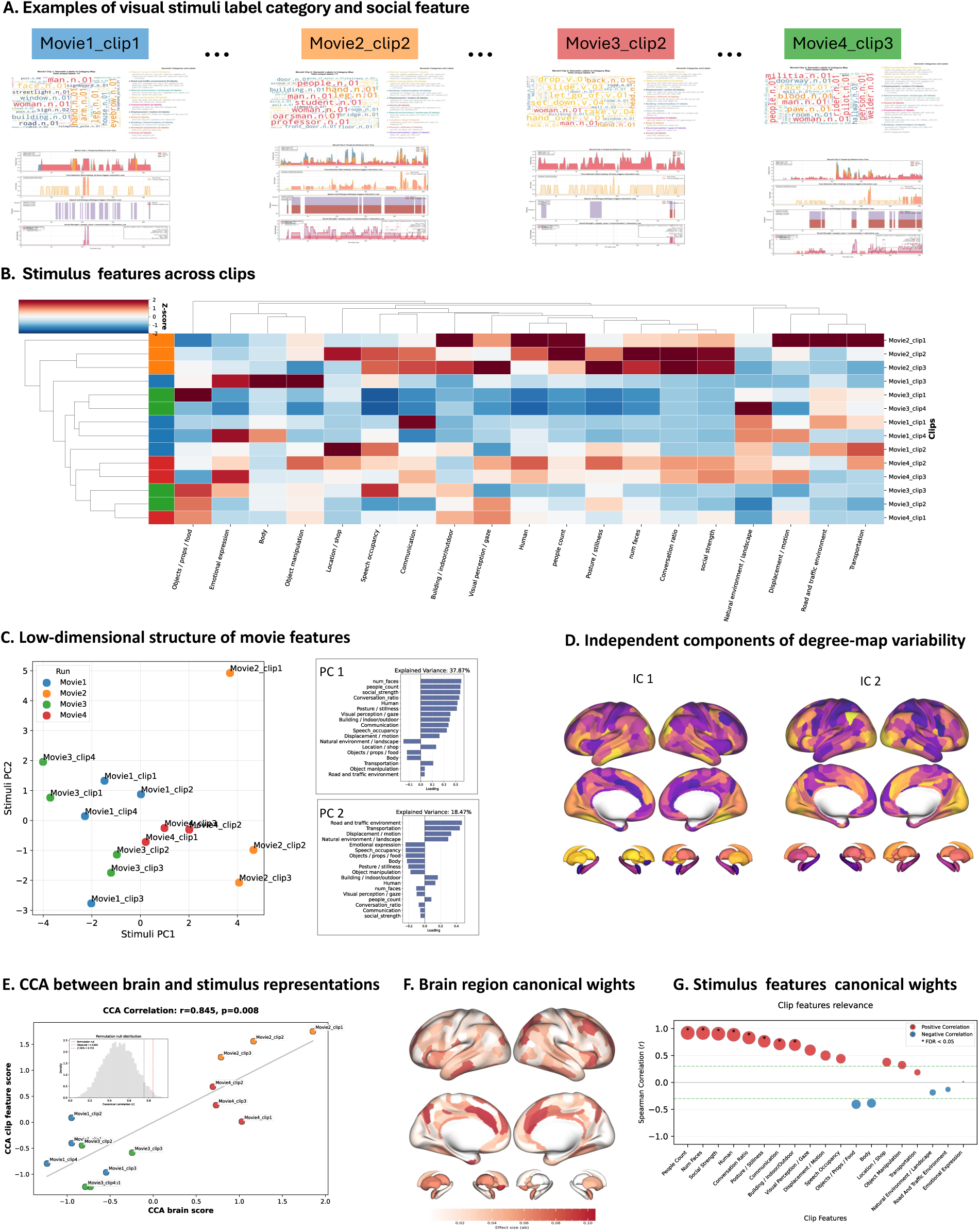
Distributed node degree-based connectivity patterns covary with latent movie feature dimensions. (A) Examples of clip-level stimulus annotations. Upper panel: Word clouds illustrate representative visual semantic label categories (WordNet-based annotations) for four example clips (Movie1_clip1, Movie2_clip2, Movie3_clip2, and Movie4_clip3). Lower panel: Time-resolved feature traces show the temporal profiles of selected stimulus dimensions aligned to the fMRI time series, including number of humans at different viewing distances (close-, medium-, and far-range faces), speech/dialogue markers, face, and a composite social-strength index. (B) Heatmap of clip-level stimulus feature intensities across the 14 clips (excluding the validation clip). Features were z-score transformed across clips and hierarchically clustered to reveal latent structure in naturalistic content, highlighting systematic differences between clips in visual, auditory, and social–semantic dimensions. (C) Low-dimensional structure of movie stimulus features across the 14 clips. Clip-level visual semantic and audiovisual human-related features were reduced using principal component analysis (PCA). The 2D scatter plot shows clips embedded in the space of the first two principal components (PC1–PC2), and bar plots indicate the feature loadings for each component (explained variance shown). (D) Independent components of node degree-map variability. Clip-level node degree maps (14 clips × 374 ROIs) were decomposed using independent component analysis (ICA), yielding two reproducible spatial components (IC1–IC2). Cortical surface maps and subcortical renderings depict the spatial loadings of each component, representing principal axes of clip-dependent node degree reweighting. (E) Canonical correlation analysis (CCA) between brain and stimulus representations. CCA was performed between ICA-derived brain scores and PCA-derived stimulus scores across clips. The scatter plot shows clip-wise canonical scores for the first canonical mode, with the observed brain–stimulus correlation and permutation-based significance (inset: null distribution). (F) Brain region canonical weights. Cortical and subcortical maps show the spatial distribution of canonical weights for the first brain canonical variate, highlighting regions contributing most strongly to the stimulus-linked pattern of node degree reweighting. (G) Stimulus features canonical weights. Contributions of individual stimulus features to the first stimulus canonical variate were quantified as correlations between each feature score and the CCA-derived stimulus canonical scores across clips. Positive and negative values indicate features loading on opposite ends of the dominant latent stimulus dimension.

To test this stimulus-brain association, we first reduced clip-level stumulus features using principal component analysis (PCA), retaining the first two components as a low-dimensional representation of stimulus features (14 clips × 2 components; Fig. 4C). In parallel, we decomposed clip-level node degree maps (14 clips × 374 ROIs) using independent component analysis (ICA), yielding two reproducible spatial components and corresponding clip-level brain scores (14 × 2; Fig. 4D). Restricting both brain and stimulus representations to two dimensions provided a parsimonious model relative to the limited number of clips. Using canonical correlation analysis (CCA), we next assessed multivariate associations between these ICA-derived brain scores and the low-dimensional stimuli features of movie content. CCA revealed a significant stimulus-brain association, with the first canonical mode showing a high correlation between brain and stimulus feature scores (Pearson r = 0.845, permutation P < 0.05; 10,000 permutations; Fig. 4E). This indicates that clip-dependent variations in node degree centrality, reflecting the total connectivity of each region with the rest of the brain, were aligned with a latent dimension of stimulus features. To assess robustness, we performed leave-one-clip-out cross-validation (LOCO-CV), in which the CCA model was trained on all but one clip and used to predict the canonical scores of the held-out clip. The predicted brain and stimulus canonical scores remained positively correlated across folds (r = 0.466), indicating that the association generalizes to unseen clips. The canonical brain pattern was dominated by dorsolateral prefrontal cortex (DLPFC), medial prefrontal cortex (MPF), anterior cingulate cortex (ACC), precuneus, inferior temporal gyrus (ITG), and superior parietal lobe (SPL) and the intraparietal sulcus (IPS) in the brain (Fig. 4F). To interpret the stimulus dimension identified by CCA, we correlated individual semantic features with the canonical stimulus scores across clips, with significance assessed using permutation testing and FDR correction. Human- and social-related features, including People Count, Num Faces, Social Strength, Human, Conversation Ratio, and Communication, together with Building/Indoor–Outdoor features, showed significant positive correlations with the canonical axis (permutation p < 0.05, FDR-corrected; Fig. 4G and Supplementary Fig. 8). In contrast, object- and location-related features showed weaker and oppositely signed associations, suggesting that object-centric scenes occupy the opposing end of the latent stimulus dimension.

### Rich-club systems in large-scale networks

Building on this conserved node degree-based backbone, we estimated which regions act as integrative hubs in movie watching and whether hub organization differs across stimuli. For each movie clip, we computed the node degree–based rich-club architecture from the group-averaged ISFC matrix. Rich-club organization refers to the tendency of high-degree nodes to form densely interconnected cores that facilitate global communication within brain networks networks ^38^. In our study, rich-club regions were defined as nodes whose degree exceeded the k value at which the normalized rich-club coefficient (ϕ_norm(k)) reached its maximum within the statistically significant range (ϕ_norm(k) > 1, p < 0.001, permutation test, Bonferroni corrected; Supplementary Fig. 9).

Across all clips, rich-club hubs showed clip-specific recruitment of subregions within higher-order perceptual and association cortex, most consistently involving posterior temporal, occipital, and parietal regions. As shown in Fig. 5, rich-club regions were consistently identified in bilateral motion-sensitive area (MT+), STG, inferior parietal lobule (IPL), superior parietal lobule (SPL), DVS and VVS, and TPOJ, with additional involvement of frontal lobe, cingulate cortex, and motor areas in a subset of clips (Fig. 5A–B). Consistent with their proposed integrative role, rich-club regions exhibited high participation coefficients across clips, indicating a predominant connector-hub profile rather than provincial connectivity (Supplementary Fig. 10). Importantly, the specific set of regions identified as rich-club hubs differed across movie clips, with partially overlapping but non-identical combinations of regions engaged in each clip (Fig.5C). Although the specific regional combinations varied across clips, rich-club organization consistently centered on a predominantly posterior set of perceptual and multimodal integrative regions (e.g., IPL, SPL, TPOJ).

**Fig. 5.**
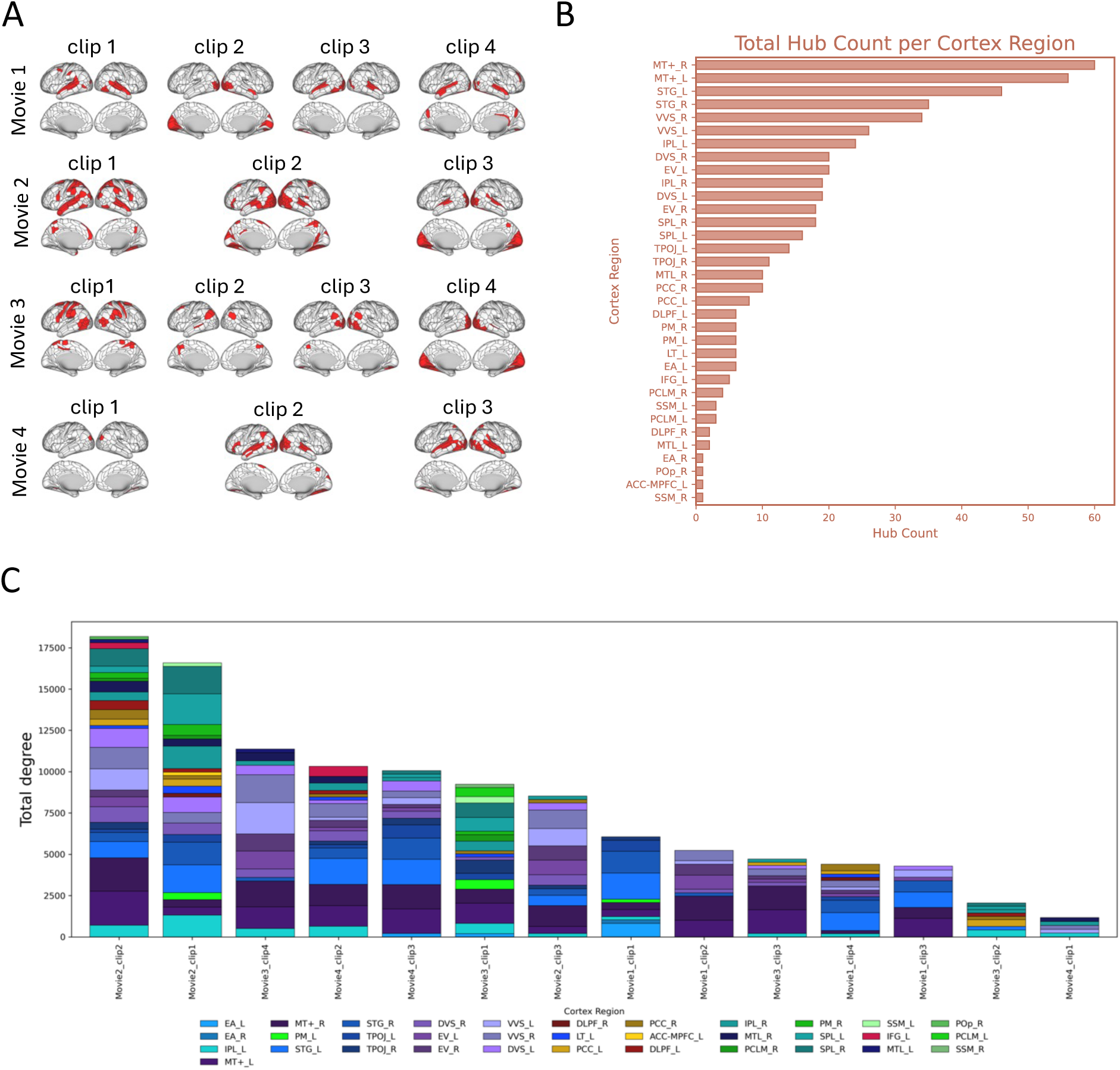
Flexible recruitment of rich-club hubs during movie watching. (A) Rich-club hub regions (marked in red) identified for each movie clip based on node degree-based rich-club analysis of group-averaged inter-subject functional correlation (ISFC) networks *(p < 0.001, Bonferroni corrected).* (B) Rich-club hubs consistency across cortical regions. Bar plot shows the cumulative hub count across clips for each cortical system/region, computed as the sum of rich-club subregions identified within each cortical eara, highlighting systems that most consistently contributed to rich-club hubs during movie watching (e.g., MT+, STG, and posterior association cortex). (C) Clip-wise rich-club composition and stimuli feature scores. Stacked bars depict the total node degree contributed by rich-club regions for each clip, partitioned by cortical regions (color-coded).

### Rich-club systems show selective connection to large-scale functional networks

To further elucidate how rich-club hubs interact with the rest of the brain during movie watching, we grouped rich-club regions into four functional systems, i.e., visual, speech, motor, and domain-general systems, based on anatomy and prior functional annotations ^36^. We then characterized their whole-brain connectivity profiles to assess how different rich-club regions mediated stimuli information to distributed cortical networks (Fig. 6B; Supplementary Fig. 11). Speech-related rich-club hubs showed positive connectivity with language areas and anterior domain-general areas (e.g., the DLPFC and posterior cingulate cortex (PCC)) and negative connectivity with visual cortex. In contrast, visual rich-club hubs showed positive connectivity to visual and posterior domain-general regions (e.g., IPL/SPL and MTL) and negative connectivity to language areas. Motor rich-club hubs showed positive connectivity to motor and visual regions and negative connectivity with language areas. Domain-general rich-club hubs exhibited a functional dissociation: parietal hubs showed preferential positive connectivity with visual areas, whereas PCC subregions preferentially positively correlated with language areas, suggesting system-specific rich-hub organization during naturalistic processing.

**Fig. 6.**
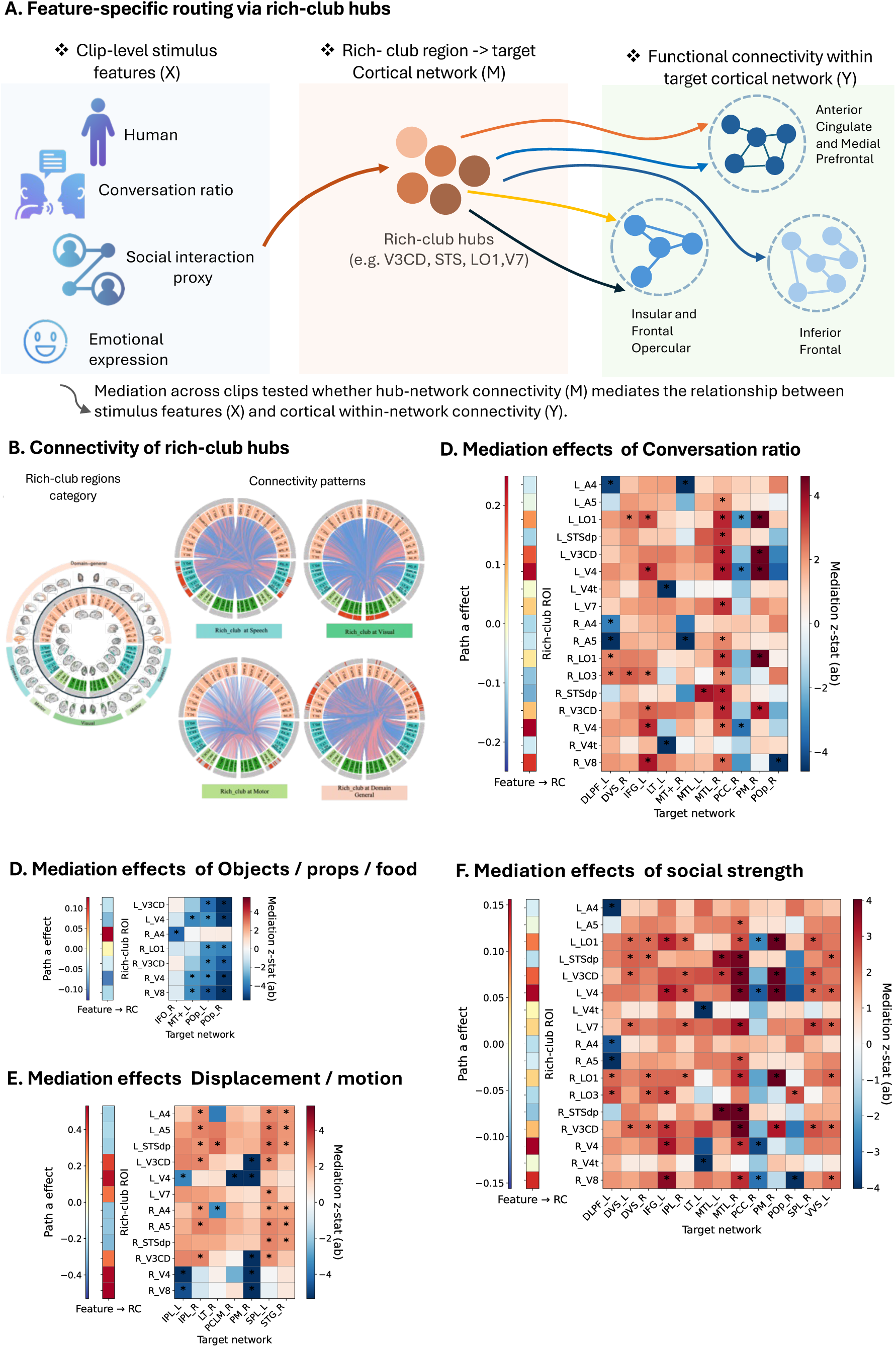
Rich-club pathways differentially mediate stimulus features to cortical networks during movie watching. (A) Conceptual schematic of the mediation framework illustrating the indirect pathway whereby clip-level stimulus features (X) influence within-network functional integration (Y) via rich-club hub–to–target network connectivity (M). Within-network integration was defined as the mean functional connectivity among all subregions within each cortical target system (e.g., anterior cingulate and medial prefrontal cortex, ACC–mPFC; inferior frontal gyrus, IFG). (B) Whole-brain connectivity profiles of rich-club hubs. Rich-club regions were grouped into four functional systems (visual, speech, motor, and domain-general; left). Circular connectivity plots (right) summarize preferential coupling patterns between each rich-club system and distributed cortical targets. (C–F) Feature-specific mediation effects. Heatmaps show permutation-based *z*-statistics of the indirect mediation effect (*a × b* path), quantifying the extent to which rich-club hub–to–target network connectivity mediates associations between stimulus features and within-network integration (a path: effect of stimulus feature on hub–to-target connectivity; b path: effect of hub connectivity on within-network integration). Adjacent bars indicate the corresponding feature-to-hub coupling effects (a path) across rich-club regions. * denote significant pathways after permutation-based testing with FDR correction (permutation p < 0.05, FDR-corrected).

### Rich-club pathways differentially mediate stimulus features to cortical networks

To test how clip-specific stimulus features is linked to network integration through rich-club hub system, we performed clip-level mediation analyses in which quantified stimulus features for each clip (X) predicted cortical within-network functional integration (Y) via the rich-club hub-to-target network connection strength (M) (Fig. 6A; Methods). For each stimulus feature, permutation-based mediation analyses identified multiple significant rich-club hubs–to–target network connectivity pathways mediating the association between stimulus features and distinct cortical networks (permutation p < 0.05, FDR-corrected; Fig. 6; Methods). For interpretability, we focused on rich-club regions that were consistently identified in at least six clips. Different stimulus features exhibited distinct mediation profiles, with each feature showing significant mediation effects linking it to multiple cortical networks through partially overlapping sets of rich-club connections (full feature set in Supplementary Fig. 12). For example, object-related content (objects/props/food) showed significant negative mediation effects, primarily involving connections between rich-club hubs and a subset of posterior–medial and temporal networks, including the fronto-opercular (FOP), posterior opercular (POP), and medial temporal lobe systems (Fig. 6D). In contrast, social- and language-related features showed broader and more distributed mediation profiles. The conversational proportion and social strength exhibited widespread significant mediation effects across multiple cortical networks via connectivity between rich-club hubs and their target networks (Fig. 6 C, F); these networks primarily involve PCC, dorsal visual regions, left IFG, IPL, and lateral and medial temporal cortex (LT/MTL). Displacement- and motion-related stimulus features showed selective mediation effects primarily involving connectivity between rich-club hubs and the inferior and superior parietal lobules (IPL and SPL; Fig. 6E), with significant hub–parietal pathways linking motion-related features to specific target network integration patterns (permutation p < 0.05, FDR-corrected). These results indicate that associations between stimulus features and within-network integration were partially mediated by feature-dependent functional connections centered on rich-club hubs across distributed cortical systems.

### Connectome topological feature verification using an in-house dataset

To validate the connectome topology observed in the HCP dataset, we analyzed an in-house movie-watching fMRI dataset with a stimulus of a 7-minute movie clip ^15^ using the same network analysis pipeline. Higher node degree and clustering were observed in STG, MTG, parietal, and occipital cortices (Fig. 7B). Rich-club organization involved bilateral temporal regions, TOPJ, ventral and dorsal visual streams, SPL, parahippocampus, and PCC (Fig. 7C). Connectivity profiling showed language-related rich-club regions preferentially positively connected with temporal and frontal areas, whereas visual and parietal rich-club regions showed stronger within-network positive connectivity and negative connectivity with language area (Fig. 7D).

**Fig 7.**
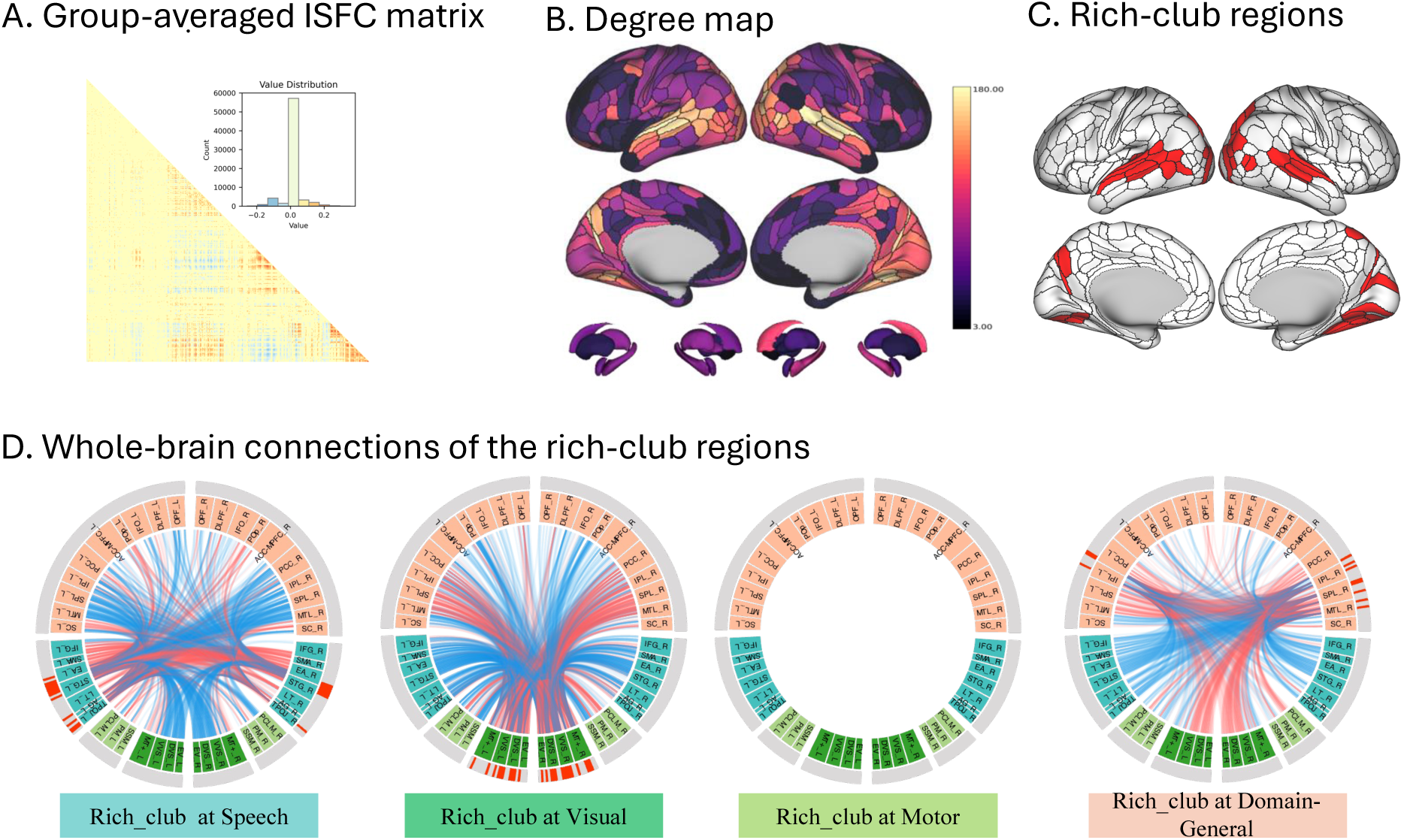
Rich-club organization in a 7-min movie-watching fMRI dataset. (A) The group-averaged ISFC matrix after removing false-positive connections through a t-test (p < 0.01, FDR corrected). (B) Node degree map. (C) Rich-club regions that have the highest normalized rich-club coefficient ⌽norm (p < 0.001, Bonferroni corrected). (D) Whole-brain connections of the rich-club regions.

## Discussion

This study reveals a dual-architecture principle underlying large-scale brain connectomic organization during naturalistic movie watching, in which stability and stimulus flexibility coexist. Specifically, movie watching consistently engages a conserved, regional node degree-based network backbone, primarily anchored in temporal and lateral occipital sensory and adjacent parietal association cortices with persistently elevated node degree, while higher-order regions engagement and rich-club hubs flexibly reconfigure in response to stimulus content. Rather than reflecting whole-brain remodeling of network topology, clip-to-clip differences primarily reflected selective reweighting of integrative participation by association regions within this conserved backbone, tightly coupled to latent dimensions of stimulus content. Importantly, rich-club analysis identified a posterior-dominant set of integrative hubs that mediate stimulus-dependent interactions between stimulus features and cortical systems via selective hub–network connectivity. Together, these findings support a mechanistic framework in which naturalistic cognition relies on a conserved poster perceptual backbone of temporo-parieto-occipital lobe, upon which stimulus-dependent processing is implemented through stimulus-dependent reweighting of regional integration strength, higher-order region engagement, and reconfiguration of rich-club organization across clips that flexibly tunes integration across large-scale functional networks.

At the global level, network topology of global efficiency and modularity followed consistent trends across density thresholds, consistent with the principle of economical small-world organization that balances integration and segregation at a relatively low wiring cost ^38^. Nevertheless, substantial differences in these metrics across clips under identical thresholds indicate that stimulus content modulates connectivity weighting within a constrained topology. This aligns with prior work showing partial cross-movie similarity in functional connectivity and activity alongside stimulus-specific patterns ^13,19,35,39^.

At the regional level, while individual regions exhibited substantial variability in absolute node degree, the overall distribution of node degree across regions remained highly similar across clips. High node degree regions consistently concentrated in lateral temporal, lateral occipital, and posterior parietal cortices, exhibited relatively small differences in degree values across clips. These perceptual and adjacent association regions play central roles in audiovisual decoding, semantic processing, and multisensory integration during naturalistic stimulation ^40–42^. Their stability likely reflects strong anatomical constraints and shared stimulus-driven synchronization, consistent with extensive neurocinematics evidence demonstrating robust inter-subject correlation in auditory and visual cortices during movie watching ^8,20,21^. Their consistent engagement across diverse movie stimuli ^22^ likely underlies the conserved network backbone we observed. In contrast, medial/orbitofrontal PFC, ACC, para-hippocampal gyrus, and subcortical regions implicated in higher-order coggition processing processing ^43^ showed systematically lower node degree and greater variability across clips. These findings provide connectome-level support for prior work showing that higher-order association systems flexibly adjust their engagement depending on naturalistic stimulus demands ^20,22,30,35,44^.

Given the clip-dependent variability in node degree superimposed on a conserved backbone, we tested how this reweighting systematically maps onto stimulus features. CCA between ICA-derived brain node degree score maps and PCA-reduced stimulus features revealed robust brain–stimulus alignment, indicating that backbone selective reweighting is not arbitrary but tracks meaningful dimensions of naturalistic content. This finding extends models of large-scale network organization that emphasize stable architectural constraints alongside context-dependent reconfiguration ^45^ into an ecologically valid naturalistic domain ^11^. The dominant brain regions included dmPFC, ACC, precuneus, and DLPFC, strongly implicated in self-referential processing, affective regulation, emotional and socioal processing ^46–48^. In addition, the involvement of ITG and parietal visual regions is consistent with their established roles in complex visual processing and goal-oriented attention, scene integration, and multi-information dimension selection attentional control during naturalistic social cognition perception ^49^. Consistent with this interpretation, our stimulus–brain CCA analysis revealed that the latent stimulus axis was strongly associated with human interaction and social related features, including people count, number of faces, social strength, coversation ratio, and communication. These findings suggest that, within the present stimulus set, the flexible engagement of higher-order association regions is preferentially driven by socially rich and human-related content in naturalistic scenes.

Taken together, our findings demonstrate that during naturalistic movie watching, clip-specific stimulus features selectively reweight the strength of regional integration within a conserved posterior perceptual–association backbone, rather than altering the global organization of the whole-brain connectome. This pattern supports a dual-architecture principle of naturalistic brain responses, whereby posterior perceptual–association systems maintain a conserved node-degree hierarchy across clips, while higher-order association systems flexibly adjust their integration in response to narrative, emotional, and social demands.

Extending the dual-architecture framework, rich-club analysis identified a posterior-dominant rich-club organization that provides a mechanistic substrate for stimulus-dependent reweighting of large-scale brain networks. Rich-club regions identified in this study include higher-order visual cortex (VVS / DVS and MT+), temporo-parietal association cortex (STG and TPOJ), and dorsal parietal regions (IPL/SPL), with additional contributions from midline integrative regions such as the precuneus and PCC. These posterior hubs have consistently shown high synchronization of activity across subjects during movie watching ^8,50,51^, with varying node degrees of spatial extent across different movie types ^19,22^.

In line with this finding, our study also found that across the 14 movie clips, the distribution and strength of those rich-club regions varied markedly, as indexed by clip-wise total node degree aggregated across cortical regions. Specifically, clips characterized by stronger social content, extended communication, and more on-screen characters, as well as those with richer scene contexts, elicited broader spatial distribution and higher overall connectivity strength of rich-club regions. Clips rich in speech or dialogue preferentially recruited STG and TPOJ as rich-club hubs, highlighting their role as integrative nodes for spoken language processing and social-cue integration during naturalistic communication ^52^. Clips featuring complex spatial transitions additionally recruited parietal and motor regions that were involved in spatial updating and intention inference ^53,54^, and in some cases recruited higher-order areas such as left IFG and DLPFC. Our results show that rich-club composition varies with stimulus demands, revealing how rich-club hub regions flexibly adjust their spatial distribution and connectivity strength on top of a conserved network backbone to support diverse stimuli feature processing during movie watching. On the one hand, our findings extend prior work on stimulus-specific activation patterns across different movies ^22^. On the other hand, whereas previous studies have primarily characterized dynamic reconfiguration of large-scale networks over short timescales within individual movies ^28,30,55^, our results show that such reconfiguration also operates across clips with distinct stimulus profiles, reflected in shifts in rich-club hub composition and hub–network connectivity.

The prominence of posterior backbone regions along this canonical axis suggests that rich-club hubs may serve as the architectural substrate through which stimulus-specific information is selectively routed across distributed systems. Mapping rich-club hubs onto language, visual, motor, and domain-general systems revealed a structured pattern of integration and segregation: speech-related hubs showed preferential coupling within language and domain-general networks, whereas visual-related hubs were more strongly embedded within visual–parietal systems, alongside reduced between-network connectivity. This organization is consistent with canonical principles of functional segregation and integration in the human brain ^56–58^. Together, these findings indicate that rich-club hubs preserve a backbone architecture while enabling content-specific routing of speech and visual information, providing a concrete network-level mechanism for balancing stability and flexibility during naturalistic cognition.

Further, individual-level mediation analysis showed that rich-club hubs mediate the association between stimulus features and integration strength within individual cortical networks, through hub-to-target connectivity from temporal, parietal, and higher-order visual regions. Each hub mediated distinct stimulus features via specialized rich-club hub–to–target network pathways, consistent with prior evidence showing that posterior temporal and parietal regions support multisensory audiovisual integration ^42^ and semantic and narrative processing during movie watching ^22^. The observation that individual stimulus features are mediated through multiple hubs–to–target network pathways toward different cortical networks supports the view that naturalistic processing relies on coordinated interactions across distributed brain systems ^59^

In particular, human-related features were associated with multiple higher-order cortical systems, with these distributed relationships being strongly mediated by connectivity between rich-club hubs and those systems. Conversational ratio showed significant mediation effects across DLPFC, IFG, DVS, MTL, and PM networks, while social interaction features additionally engaged POP and VVS systems. These networks support language, social inference, executive control, memory integration, and event representation critical for conversational and social processing ^22,60–63^. These findings suggest that, under naturalistic conditions, the integration of socially and human-related information indeed relies on coordinated processing across multiple higher-order cortical systems, mediated by rich-club connectivity that facilitates cross-system coordination during complex social interpretation. Displacement- and motion-related stimulus features showed selective mediation effects primarily involving connectivity with the IPL and SPL, in line with the established role of these regions as core nodes of the dorsal perceptual–action stream supporting spatial transformation and motion tracking ^5,64,65^. In contrast, object features showed predominantly negative rich-club mediation to IFO, POP, and MT+, consistent with prior work showing object recognition relies on specialized ventral visual and semantic pathways ^66,67^ that reduce the dependence for large-scale network coordination.

The presence of multiple hub-mediated pathways linking individual stimulus features to diverse cortical networks supports the view that rich-club hubs function as coordination nodes that enable stimulus-dependent modulation of network integration without disrupting global topology. Accordingly, because different movie clips emphasize distinct stimulus features, they are likely to engage partially different configurations of rich-club hubs to support content-specific processing demands.

To test the generalizability of the connectome backbone, we repeated the degree and rich-club analyses in our inhouse movie-watching fMRI dataset with a single continuous 7-min movie clip as the stimulus. This longer stimulus minimizes potential carryover effects across clips and provides a complementary test of topological stability under sustained naturalistic viewing. We observed a highly convergent configuration that closely matched the backbone identified in the HCP dataset. This cross-dataset replication supports the robustness of temporo-occipital and parietal association regions as a stable integrative backbone during naturalistic movie watching.

Guided by the hypothesis that large-scale functional topology is stimulus dependent, we further discussed which types of movie content most reliably engage the connectome backbone across individuals. Our results showed that Hollywood films did not uniformly outperform other stimuli. Specifically, characteristics and rich-club organization in independent films were comparable, and one Hollywood clip (Movie 4, clip 1) exhibited weaker organization than an independent film clip. Nevertheless, several Hollywood clips (e.g., Movies 2 and 3, clips 2–3) showed high pairwise similarity, likely reflecting shared narrative structure, interpersonal focus, and a clear central character ^22,68^. Consistent with prior work showing that abstract, nonverbal stimuli resemble resting-state connectivity responses ^69^, clips lacking dialogue or narrative structure elicited weaker topology and predominantly visual organization. In contrast, socially and narratively rich clips, across both Hollywood and independent films, produced the strongest and most reliable rich-club recruitment, particularly within temporal and parietal and higher-level visual systems. Together, these findings indicate that stimuli rich in socially and behaviorally meaningful content most reliably elicit stable and reproducible backbone configurations across individuals, imposing a content-dependent constraint on the stability of large-scale network topology during naturalistic movie watching.

Several limitations should be noted in this study. First, the number of clips was modest relative to the dimensionality of semantic features, which may bias multivariate alignment toward the dominant scene–object axis. Larger stimulus sets will be needed to more fully dissociate semantic dimensions, including social and affective factors. Second, although we replicated the degree and rich-club backbone in an inhouse dataset with a longer and continuous stimulus, this in-house dataset did not include matched semantic feature annotations, preventing an external validation of the feature-alignment and mediation analyses. Future work combining longer continuous stimuli with time-resolved semantic labeling will be essential for testing whether the same stimulus dimensions drive backbone reweighting across datasets. Third, mediation analyses quantify statistical indirect pathways but do not establish causal directionality, motivating future work using effective connectivity models or perturbational approaches. Fourth, because rich-club hubs were defined from node degree-based topology under thresholded connectivity, future analyses should evaluate parameter sensitivity and extend to weighted or multilayer formulations. Despite these limitations, the convergence across clips and with an independent dataset supports a generalizable backbone architecture that is flexibly reweighted in a content-dependent manner.

## Conclusion

The findings of this study demonstrate that naturalistic cognition is supported by a dual-architecture principle of large-scale brain networks: a converged, posterior node degree-based backbone ensures reliable audiovisual integration, while higher-order association regions flexibly reweight their participation according to stimulus content. Variability within this backbone is systematically aligned with socially and behaviorally salient features, and is implemented through rich-club–mediated across functional systems. Together, these findings provide a mechanistic account of how stability and flexibility are balanced in the human connectome during movie watching.

## Methods

### Subjects and experimental paradigm of the Human Connectome Project 7T dataset

The Human Connectome Project (HCP) Young Adult 7T release dataset 47 was used in this study from the original 184 subjects, data of 176 (106 females and 70 males) subjects who underwent all four movie-watching runs (Movies 1-4) were selected for this study. All subjects were healthy young adults aged between 22 and 35 years, and all scanning sessions were conducted at the Center for Magnetic Resonance Research at the University of Minnesota. For this HCP dataset, each movie run consisted of 4 or 5 clips separated by 20 seconds of rest blocks. Movie 1 and Movie 3 featured clips from independent films (both fiction and documentary films). The other two runs, Movie 2 and Movie 4, included clips from Hollywood films. For a brief description of each clip, see ***Supplementary Table 1***. All movie clips were sourced from the Human Connectome Project (HCP; https://db.humanconnectome.org/).

Movie-watching fMRI data were acquired on a 7.0 Tesla Siemens Magnetom scanner Functional blood oxygen level-dependent (BOLD) images were acquired in a single-shot gradient-echo echo-planar imaging (EPI) with a Nova32 32-channel head coil: TR = 1000 ms, TE = 22.2 ms, flip angle = 45°, field of view (FOV) = 208 × 208 mm, in-plane resolution = 130 × 130, voxel size = 1.6 mm isotropic, 85 axial slices, multiband acceleration factor = 5, in-plane acceleration (iPAT) = 2, partial Fourier = 7/8, echo spacing = 0.64 ms, and bandwidth = 1924 Hz/Px. fMRI data were acquired with alternating phase-encoding directions: Movie 1 and Movie 3 in the anterior–posterior (AP) direction, and Movie 2 and Movie 4 in the posterior–anterior (PA) direction. The total durations of the four movie runs were 921 sec, 918 sec, 915 sec, and 901 sec, respectively.

### Subjects and experimental paradigm of the in-house dataset

We used the movie-watching fMRI data that was reported in Tie, et al.^15^ for this study. This dataset included movie-watching fMRI data from 22 right-handed healthy subjects (11 females and 11 males, mean age = 26.3 years, range: 19-39 years) who had no history of neurological, cognitive, or psychiatric disorders, nor any speech, hearing, or vision deficits. All subjects were native English speakers. The study protocol was approved by the Mass General Brigham Institutional Review Board, and all subjects provided written informed consent. For this dataset, the movie stimulus was a 7-minute excerpt from the family film *The Parent Trap* (1998; directed by Nancy Meyers; produced by The Meyers/Shyer Company and Walt Disney Pictures).

Movie-watching fMRI data were acquired on a 3.0 Tesla GE Signa system (General Electric, Milwaukee, WI, USA). fMRI images were acquired in a single-shot gradient-echo EPI and a standard quadrature head coil: TR= 2000 ms, TE = 40 ms, flip angle = 90°, slice gap = 0 mm, FOV = 256 × 256 mm, in-plane resolution = 128 × 128, voxel size = 2 × 2 × 4 mm^3^, 27 axial slices, ascending interleaved sequence. Structural images were acquired using a T1-weighted 3D spoiled gradient recalled (SPGR) spatial sensitivity encoding technique (ASSET, i.e., parallel imaging) and an 8-channel head coil: TR= 7.8 ms, TE = 3.0 ms, flip angle = 20°, in-plane resolution = 512 × 512, voxel size = 0.5 × 0.5 × 1 mm^3^, 176 axial slices.

### Data analysis

#### Preprocessing

The HCP movie-watching fMRI data were minimally preprocessed using the minimal HCP pipeline of fMRIVolume ^71,72^. The first 10-sec data of all fMRI runs was discarded to allow for stabilization of the BOLD signal. Preprocessing included gradient distortion correction, TOPUP-based spin echo field map unwarping, motion correction, high-pass filtering with a cutoff of 2000 seconds, brain-boundary-based registration of EPI to structural T1 images, non-linear registration to Montreal Neurological Institute (MNI) standard space and grand-mean intensity normalization. fMRI data were cleaned up for head motion, cardiac pulsation, breathing, and scanner artifacts using sICA+FIX denoising. In addition, the average whole-brain signal at each TR was calculated using an HCP-provided per-subject, per-run mask (brainmask_fs.1.60.nii.gz) and regressed out from the FIX-denoised images. The data were then spatially smoothed using a 4 mm Full Width at Half Maximum (FWHM) Gaussian kernel and interpolated to 2 mm isotropic voxels.

To account for the hemodynamic delay, we included 5 TRs (5 seconds) following the ending of each clip in the functional connectivity calculation, and the remaining 15 TRs (15 seconds) of the 20-second rest period between clips were excluded from the analysis.

The in-house dataset movie-watching fMRI data were preprocessed using the pipeline developed by the CBIG group (https://github.com/ThomasYeoLab/CBIG/tree/master/stable_projects/preprocessing/CBIG_fMRI_Preproc2016). The first 10-sec data of fMRI was discarded. Then Motion correction was performed using spline interpolation, and Framewise Displacement (FD) and DVARS were calculated. Frames with excessive motion (FD > 0.2) or signal changes (DVARS > 50) were regarded as outliers. The corrected EPI images were registered to individual structural T1-weighted images and subsequently normalized to the 2 mm MNI152 standard space. fMRI data were denoised by regressing out several confounding signals, including the global mean signal, white matter (WM) and cerebrospinal fluid (CSF) signals, as well as 12 motion-related regressors (6 original parameters and their 6 temporal derivatives). Band-pass filtering (0.01–0.1 Hz) was then applied using a regression approach to capture neural signals relevant to both rest and task states, followed by spatial smoothing with a 4-mm full-width at half maximum (FWHM) Gaussian kernel.

### Brain network construction

The brain functional network were constructed on 374 regions of interest (ROIs) including 360 cortical ROIs (180 in each hemisphere) defined by the HCP MMP 1.0 atlas 25 and 14 subcortical ROIs (7 in each hemisphere) defined by the Harvard-Oxford atlas (https://fsl.fmrib.ox.ac.uk/fsl/docs/#/other/datasets). The preprocessed fMRI signals from all voxels within each ROI were extracted and averaged to generate a regional mean time series, which was subsequently Z-transformed across time points to normalize signal amplitude. This yielded a single representative time course per ROI, which was then used for subsequent functional connectivity analyses. For each subject, the weighted brain network connections were constructed via a leave-one-out inter-subject functional correlation (ISFC) approach 36. ISFC is a powerful method for studying brain function during task states, offering improved signal-to-noise ratio (SNR) and a more comprehensive view of functional integration compared to traditional intra-subject network analyses, such as temporal correlations (Pearson’s *r*) between the time series of each parcel pair are computed within individual brains ^30,32^. For each subject, the ISFC matrix shows the functional connectivity of each ROI calculated by the Pearson correlation between the time course of that subject and the average time course of all other ROIs across all other subjects. Pearson correlation coefficients were then transformed to z-scores using Fisher’s transformation. The ISFC connectivity matrix was thresholded using a one-sample t-test: p < 0.01 (Bonferroni correction) for the HCP dataset) and p < 0.01 (FDR correction) for the in-house dataset. The connections with correlation coefficients (r) exceeding the threshold were considered statistically significant and retained in the final ISFC matrix. Finally, a symmetry group-based ISFC matrix was computed by averaging individual subjects’ connectivity matrices for further analyses.

### Graph metrics calculation

We used graph-theoretical network measures to characterize the topology of the functional connectome ^73^ in this study. For a given network matrix, each brain ROI was defined as a node and the connectivity between ROIs as an edge. Then the graph metrics of each mean-ISFC of each clip was computed using the Brain Connectivity Toolbox for Python (https://brainconn.readthedocs.io/). At the node level, degree was computed to quantify regional connectivity strength. At the global level, we assessed network efficiency and modularity to characterize large-scale integration and segregation. Degree maps were used as the primary regional metric for subsequent multivariate and mediation analyses. We further quantified rich-club organization ^26^, a property of complex networks in which a subset of highly connected nodes (hubs) are more densely interconnected and support efficient global communication. Statistical significance was assessed by comparison with degree-preserving randomized networks (1,000 permutations), yielding normalized rich-club coefficients. Rich-club regions were defined as nodes with degree exceeding the threshold that maximized the normalized rich-club coefficient for each clip. Across clips, we further assessed the centrality profile of rich-club regions using participation coefficients to distinguish connector versus provincial hubs. Detailed definitions, formulas, and statistical procedures for graph metrics and rich-club analysis are provided in the Supplementary Methods. Building on this framework, we identified hub regions involved in processing the diverse movie clip stimuli.

### Statistical analysis of between-clip variability in network topology

To evaluate whether brain network topology varied systematically across movie clips, we conducted an one-way repeated measures ANOVA with clip as the independent variable and each ROI’s graph metric (i.e., degree, clustering coefficient, and betweenness centrality) as the dependent variables. This analysis tested whether the distributions of these network measures differed significantly across clips. For post-hoc comparisons, we assessed pairwise similarities between clips by computing Pearson correlation coefficients of each graph metric’s regional distribution. In addition, we calculated the coefficient of variation (CV, defined as the standard deviation divided by the mean) for each metric across clips to quantify the relative variability of regional topology.

### Cortical-area–level and functional based framework for rich-club network categorization

To interpret rich-club core locations and enable subsequent system-level mediation analyses, we summarized rich-club organization at the level of 22 cortical regions and one subcortical category, following the ROI to cortical-area grouping annotations provided by the HCP-MMP1.0 atlas ^36^. The full names and abbreviations of each cortical area in the Supplementary Table 4. For each clip, ROI-level rich-club hubs were first identified, and hub membership was then aggregated within each cortical area to characterize cross-clip patterns of hub recruitment at a coarse anatomical scale.

To further characterize the functional architecture of rich-club regions, we divided the whole brain into four large-scale functional groups based on anatomical and functional affiliations ^36^: speech, visual, motor, and higher order domain-general systems, and rich-club regions were assigned to their respective functional group. For each movie clip, we computed the ISFC patterns of rich-club regions within each subgroup relative to the whole brain. Specifically, we extracted the ISFC profile of each rich-club region and visualized their connectivity pattern to the rest of the brain regions, highlighting differences across functional subgroups. This approach enabled us to investigate how rich-club regions, distributed across distinct functional systems, dynamically reorganized their connectivity in response to different stimulus conditions.

To quantify stimulus features, we leveraged the TR-resolved semantic annotations provided with the HCP 7T movie-watching dataset (https://balsa.wustl.edu/). These annotations label the presence of multiple stimulus features at each time point for every clip using WordNet-based categories, which were summarized for visualization using word clouds (Supplementary Fig. 2). The resulting feature set captures a set of predefined visual semantic dimensions spanning human and social signals as well as scene and object content, including: Human, Body, Face, Building/indoor/outdoor, Road and traffic environment, Transportation, Natural environment/landscape, Objects/props/food, Location/shop, Displacement/motion, Posture/stillness, Visual perception/gaze, Communication action, Emotional expression, and Object manipulation (Supplementary Fig. 2).

In addition to these visual annotations, we extracted speech- and social-related features at both TR and clip levels. Speech presence was detected using pyannote ^74^, and face-related features were extracted using Pliers ^75^. At the TR level, we binarized speech and conversation as 0/1 indicators reflecting whether speech or dialogue was present at each TR. At the clip level, we summarized these measures as speech occupancy and conversation ratio, defined as the proportion of TRs labeled as speech or conversation within the clip, respectively. To quantify human presence, we detected people in each frame using YOLO ^76^ and computed the number of individuals per TR. To reduce the influence of background bystanders (e.g., distant pedestrians) on social estimates, we restricted social computations to close-range people, defined using bounding-box size and position thresholds: a detection was classified as close-range if its bounding-box area exceeded 0.08 of the frame (close_area_min = 0.08) and its bottom coordinate was below 70% of the frame height (close_bottom_min = 0.7). Conversely, detections with bounding-box area < 0.03 (far_area_max = 0.03) and bottom coordinate < 50% of frame height (far_bottom_max = 0.5) were treated as far-range and excluded from social strength calculations. We further defined a measure of social strength as a composite index of social intensity for each ^77^. Specifically, we defined an interaction cue as a binary variable set to 1 when interaction-related WordNet labels (e.g., talk.v.01, conversation.n.01, gaze.v.01, look.v.01, argue.v.01, kiss.v.02, hug.v.01) were present at a given TR, and 0 otherwise. Social strength was then computed as:

social strength = (number of close-range people) × (interaction cue) × (communication).

The feature time courses for each clip are shown in Supplementary Fig. 2. For clip-level analyses, we summarized each stimulus dimension by averaging its values for each TR across the clip, yielding a 14 (clips) × 19 (features) matrix capturing clip-level stimulus feature profiles (see Fig 2D). To characterize the structure of this semantic feature space, we examined (i) feature-by-feature similarity across clips (Supplementary Fig. 3A), and (iii) clip-by-clip similarity of overall semantic profiles (Supplementary Fig. 3B). These analyses demonstrate that movie clips differ systematically in semantic composition and that stimulus dimensions exhibit structured covariance, providing a principled basis for subsequent dimensionality reduction and brain–stimulus association analyses.

A multivariate stimulus-brain canonical correlation analysis (CCA) between clip-level node degree maps and clip-level stimulus feature profiles was performed to test whether clip variations in node degree organization covary with stimulus featuresFirst, to extract the dominant axes of variability in node-degree organization across clips, we applied independent component analysis (ICA) ^78^ to the node-degree matrix. To ensure robustness, ICA was repeated multiple times with different random initializations, and component stability was assessed using an ICASSO framework ^79^, whereby resulting components were clustered to identify reproducible sources. Specifically, FastICA was repeated 100 times with different random initializations, and components were clustered to identify reproducible spatial centroids (Supplementary Methods). Based on stability and the limited number of observations and to avoid overfitting, we retained two independent components (ICA1–ICA2), yielding clip-level brain scores (14 × 2) as a low-dimensional representation of node degree-map variability.

In parallel, the 14 × 19 stimulus feature matrix was reduced using principal component analysis (PCA). Given the rapid drop in explained variance after the first few components and the limited number of clips (n = 14), we retained the first two PCs (PC1–PC2), which together explained 56.34 % of the variance in stimulus content (PC1: 37.87%, PC2: 18.47%). Clip-level PC scores (14 × 2) were used as a low-dimensional representation of stimulus features. Multivariate associations between ICA-derived brain node degree scores and PCA-derived stimulus features scores were quantified using canonical correlation analysis (CCA) ^80^, which identifies linear combinations of brain and stimulus variables that maximize their correlation. Prior to CCA, both score matrices were z-scored across clips. We fitted a one-component CCA model (n_components = 1) and defined the canonical correlation as the Pearson correlation between the resulting canonical variates. Statistical significance was assessed using a permutation test (10,000 permutations).

To interpret the spatial distribution of the brain canonical variate, we reconstructed an ROI-level canonical weight map by linearly combining the CCA weights on brain ICA components with the corresponding ICA spatial loadings (Supplementary Methods). Contributions of individual stimulus features were quantified as correlations between each original stimulus feature and the CCA-derived stimulus canonical scores across clips.

To quantify whether rich-club hubs mediate the relationship between movie stimulus features and large-scale network integration, we performed subject-level mediation analyses using clip-level functional connectivity data from 176 participants across 14 movie clips. This approach is essential for testing whether the effect of one variable on an outcome is partially or fully transmitted through an intermediate mediator ^81^. For each mediation model, the input (X) was each clip-level stimulus feature, the mediator (M) was defined as the mean functional connectivity between a given rich-club hub and all ROIs within a target cortical area, and the outcome (Y) was defined as the mean functional connectivity among all ROI pairs within that cortical area. Analyses were restricted to rich-club regions that were identified as rich-club members in at least 6 clips to ensure robustness across stimuli.

For each clip, subject-level adjacency matrices were derived from preprocessed ISFC networks (retaining the top 30% strongest connections; Fisher z-transformed and tanh-normalized). For each subject and clip, connectivity was summarized separately for (i) hub–target network connectivity (M; edges linking each rich-club hub to all ROIs within a target cortical region) and (ii) target network integration (Y; edges among ROIs within the target cortical region). Target cortical regions were defined using the ROI-to-region grouping annotations provided by the HCP-MMP1.0 atlas ^36^. Unless otherwise specified, connectivity measures were computed as the mean edge weight.

Clip-level Mediation analyses were conducted independently for each stimulus feature and each rich-club hub–target network pair. For each subject, indirect effects (ab) were estimated across clips, where a denotes the effect of the feature on hub–network connectivity (a: X→M) and b denotes the effect of hub–network connectivity on within-network connectivity (b: M→Y), controlling for X. Group-level indirect effects were summarized using bootstrap resampling across subjects (5,000 iterations; mean and 95% confidence intervals). Statistical significance was assessed using clip-label permutation testing (5,000 permutations), yielding permutation-based p values (pₚₑᵣₘ) and standardized z statistics (zₚₑᵣₘ). For each feature, significance of hub–network pathways was assessed with p < 0.05 FDR correction across rich-club hubs within each target network. Significant pathways were interpreted as evidence that feature–integration associations were partially mediated by specific rich-club hubs–to–target network connectivity patterns, as quantified by the indirect effect (ab).

## Acknowledgement

We thank Alexandra J. Golby, MD, for her support with the in-house data collection and the overall project. This research is supported by the National Institutes of Health (NIH) (grants R01DC020965 and R21NS075728).

## Author Contributions

X.W., E.L., and Y.T. designed the study. X.W. developed the methodology and conducted data analyses. E.L. and Y.T. provided guidance on data analysis and interpretation. L.R., C.P.G., and Y.T. contributed to in-house data collection. X.W., C.P.G., and Y.T. wrote the manuscript. All authors discussed the results and approved the final manuscript.

