## Supplementary Materials for "A conserved node degree-based backbone and flexible hub organization of brain connectome underlying naturalistic movie watching"

***Corresponding author**

### Supplemental method

#### Graph-theoretical metrics and rich-club analysis

*Network construction*

For each movie clip, the group-averaged ISFC matrix $M \in R^{N \times N}$ was thresholded at a fixed density. Then graph metrics of a given network matrix M were computed as follows.

*Degree and clustering coefficient of a node*

The degree of a node represents the number of connections (edges) it has to other nodes in the network. The degree $d_i$ of a node i is defined as:

$$d_i = \sum_{i \neq j} b_{ij}$$

Where**:**

$b_{ij}$: denotes the binary adjacency matrix, where $b_{ij} = 1$ where if a functional connection exists between nodes $i$and $j$after thresholding, and $b_{ij} = 0$ , otherwise.

*Global efficiency*

Global efficiency of the brain network refers to a measure of how effectively information or resources are exchanged across the entire brain. It is defined based on the shortest path lengths between nodes:

$$E_{global} = \frac{1}{N(N-1)} \sum_{i \neq j} \frac{1}{d_{ij}}$$

Where:

$d_{ij}$: Shortest path length (geodesic distance) between regions $i$ and $j$,

N: the total number of possible node pairs in the network.

*Modularity*

Modularity is a measure of the extent to which a network can be divided into distinct, densely interconnected groups of nodes (i.e., modules or communities) with sparse connections between groups. In this study, we examined the modular organization of brain functional networks using the Louvain algorithm [26,82].

*Rich-club organization*

We examined the rich-club organization of the brain functional networks by computing the rich-club coefficients ϕ(k) across a range of degrees k [83]. The weighted rich-club parameter $\phi^{w}(k)$ was computed as the ratio between the total weight of connections among nodes with degrees greater than k ($W_{>k}$ ) and the sum of the weights of the strongest $E_{>k}$ connections in the entire network: $\phi^{w}(k)$ is defined as:

$$\phi^{w}(k) = \frac{W_{>k}}{\sum_{i=1}^{E>K} \omega_i^{ranked}}$$

where:

$E_{>k}$ is the number of connections among nodes with degrees greater than k, and the weights are derived from the ranked connections in the network.

$W_{>k}$ is the total weight of connections among nodes with degrees greater than k,

$\sum_{i=1}^{E>K} \omega_i^{ranked}$ is the sum of the weights of the to $E_{>k}$ connections in the network, ranked by their weights.

To assess the statistical significance of the rich-club organization, a permutation testing was employed using a population of 1,000 randomized networks, preserving the degree distribution and sequence of the original network. For each degree k, the weighted rich-club coefficient ϕwrandom(k) was computed for each of the randomized networks. The average rich-club coefficient ϕwrandom(k) was then calculated as the mean value across m random networks. Finally, the normalized rich-club coefficient ϕwnorm(k) was computed as:

$$\phi^{wnorm}(k) = \frac{\phi^{w}(k)}{\phi^{wrandom}(k)}$$

A significant rich-club organization (i.e., $\phi^{norm}(k) > 1$) indicates that high-degree nodes are more interconnected than chance. Statistical significance was evaluated using Bonferroni-adjusted α-levels of 0.0001 per test (0.01 divided by the number of tests performed ; see ***Supplementary Fig. 9***).

*Rich-club centrality*

We assessed the centrality of the rich-club to evaluate its role in the global network structure. Specifically, rich-club regions were classified as "provincial hubs" if they exhibited a high within-module degree, and as "connector hubs" if they showed a participation coefficient (PC) that is higher than 0.5 [38]. The within-module degree identifies regions that are highly connected within their own functional community, whereas the PC quantifies the diversity of inter-module connections, with values above 0.5 indicating substantial cross-community integration. These measures highlight the extent to which the rich-club region serves as a backbone for both local specialization and efficient global communication across the entire brain.

**Stable ICA decomposition (Independent Component Analysis with Stability Analysis by Clustering)**

Clip-wise degree maps were arranged as a matrix $X \in R^{14\times374}$ (clips × regions) and z-scored across clips for each region. For a given number of components ($n_{IC}$), FastICA (max_iter = 100, unit-variance whitening) was repeated 100 times with different random initializations. Resulting component maps were L2-normalized and sign-aligned. All components across runs were clustered using agglomerative clustering based on absolute spatial correlation (distance = $1-|\ r\ |$). Cluster centroids defined stable components, and stability was quantified as the mean absolute correlation between individual components and their centroid. Component stability was assessed across candidate values of $n_{IC}$ = 2–6 (Supplementary Fig. S4). ICASSO-like stability analysis indicated high reproducibility for the two components (mean stability ≈ 0.99), whereas additional components showed reduced stability (Supplementary Fig. S4). Based on this stability profile and to retain a parsimonious representation of the data, $n_{IC} = 2$ was selected for subsequent analyses.

**Dimensionality reduction of semantic features**

To obtain a low-dimensional representation of movie content comparable to the brain ICA representation, we applied principal component analysis (PCA) to the clip-wise semantic feature matrix (14 clips × 18 features). Examination of the scree plot revealed the first two principal components explaining 56.34% of the variance (Supplementary Fig. S5). We therefore retained the first two components for canonical correlation analysis to capture the dominant covariance structure of semantic features.

**CCA and permutation test.**

(i) Input matrices. We loaded clip-level ICA scores (rows = clips; columns = IC1–IC2) and clip-level semantic PCA scores (rows = clips; columns = PC1–PC2). Clips were aligned by clip identifiers to ensure identical ordering across matrices.

(ii) Standardization**.** Each column of the brain matrix $G$and stimulus matrix $S$ was standardized across clips to zero mean and unit variance (z-scoring).

(iii) CCA fitting**.** We fit CCA with $n_{components} = 1$to obtain canonical variates $u = Ga$ and $v = Sb$ . The observed canonical correlation was computed as $r_{\mathrm{obs}} = \mathrm{corr}(u, v)$ .

(iv) Permutation test**.** To test $H_0$ that brain–stimulus alignment is not clip-locked, we permuted clip order on the stimulus matrix only (row permutation of $S$), refit CCA for each permuted dataset, and computed $r_{\mathrm{perm}}$. We used 10,000 permutations with a fixed random seed for reproducibility. Two-tailed p values were calculated as

$$p = \frac{\{|\ r_{\mathrm{perm}}\ |\geq|\ r_{\mathrm{obs}}\ |\} + 1}{N_{\mathrm{perm}} + 1}.$$

To assess robustness of the canonical correlation under resampling variability, we performed bootstrap resampling across clips (5,000 bootstrap samples). In each bootstrap iteration, clips were sampled with replacement, and the CCA model was refit to the resampled brain and stimulus matrices. The canonical correlation was recomputed for each resample, yielding a bootstrap distribution of correlation values. We report the mean and percentile-based 95% confidence intervals of this distribution（Supplementary Fig. S6）.

We assessed generalization using leave-one-clip-out cross-validation (LOCO-CV), where the CCA model was trained on 13 clips and evaluated on the remaining held-out clip in each fold. Predicted canonical variates from all held-out folds were concatenated and correlated to quantify out-of-sample performance. This procedure was used to test whether the model generalized beyond the training data and to evaluate potential overfitting. The model retained moderate out-of-sample performance under LOCO-CV ($r = 0.4660$；Supplementary Fig. S7), indicating a robust and generalizable association between brain and semantic representations.

**Derivation of ROI-level canonical brain weights**

Canonical weights on the brain side were first extracted from the CCA solution as the weight vector associated with the ICA-derived brain scores for the first canonical variate. To ensure scale invariance, this weight vector was normalized to unit length.

Stable ICA decomposition provided spatial loadings for each independent component across all graph-metric–by–ROI features. To reconstruct the canonical brain pattern in the original feature space, ICA loadings were linearly combined using the normalized CCA brain weights. Specifically, the canonical weight for each graph-metric–ROI feature was computed as the weighted sum of its ICA loadings across components. The Canonical brain pattern reconstruction as following:

$\mathrm{X} \in \mathrm{R}^{N \times F_b}$ denote the clip-wise brain feature matrix (graph metrics × ROIs),

$\mathrm{A} \in \mathrm{R}^{K \times F_b}$ denote the stable ICA spatial loadings (with $K$ independent components),

$\mathrm{w}_b \in \mathrm{R}^{K}$ denote the CCA weight vector associated with the brain-side canonical variate.

The canonical brain weight vector in the original feature space was reconstructed as:

$$\mathrm{c}_b = \mathrm{A}^{\mathsf{T}} \mathrm{w}_b$$

To facilitate comparison across components, the CCA weight vector was normalized to unit length prior to projection. Feature-level canonical weights were subsequently aggregated at the ROI level. The primary measure reported was the L2 norm of canonical degree weights within each ROI, reflecting the overall contribution strength of that region to the canonical brain pattern, independent of sign. Signed mean weights were additionally examined to preserve directional information. For each ROI $r$, the canonical contribution was summarized as the L2 norm of its degree-based canonical weights:

$$c_r = \sqrt{\sum_{i \in r} w_i^2}$$

where $w_i$ denotes the canonical weight associated with node degree for ROI $r$

**Stimulus feature relevance to the stimulus canonical variate**

To quantify which stimulis features were most aligned with the stimulus canonical axis, we computed spearman rank correlations between $s$and each stimulis feature vector $X_{:,j}$across clips:

$$\rho_j = \mathrm{Spearman}(\mathrm{w}_s, X_{:,j}), j = 1, \ldots, F.$$

$X \in \mathrm{R}^{N \times F}$denote the clip-wise feature matrix ( $N = 14$ clips; $F$ stimulus features), aligned to the clip order used in the CCA. Let $\mathrm{w}_s$, denote the stimulus-side canonical score (canonical variate) for the first CCA dimension.

Two-sided p-values were obtained for each correlation and corrected for multiple comparisons across features using the FDR procedure. Features were ranked by $| \rho_j |$, and we report features exceeding a predefined effect-size threshold (e.g., $| \rho |> 0.3$) together with their FDR-adjusted significance.

### Mediation anylysis

We performed subject-level mediation analyses to test whether rich-club hub connectivity mediates the relationship between movie semantic features and large-scale network integration. Analyses were based on clip-wise functional connectivity from 176 participants across 14 movie clips and were restricted to regions identified as rich-club members in at least 6 clips. Mediation analyses were conducted at the subject level to preserve inter-individual variability in functional connectivity and to avoid inflating statistical evidence through clip-level averaging.

For each subject $s$, clip $c$, rich-club hub $r$, and target sub-netwoek $l$, the mediator and outcome variables were defined as:

$$M_{s,c}^{(r,l)} = \frac{1}{\mid l \mid}\sum_{i\in l} \mid W_{s,c}(r,i) \mid$$

$$Y_{s,c}^{(l)} = \frac{2}{\mid l \mid (\mid l \mid -1)} \sum_{i<j,\, i,j\in l} \mid W_{s,c}(i,j) \mid$$

where $W_{s,c}$denotes the subject-level functional connectivity matrix (top 30% density, Fisher *z*-transformed, tanh-normalized). Analyses focused on absolute edge weights; signed variants yielded comparable results.

For each subject, mediation was estimated across clips using linear models:

$$M_{s,c} = a_s X_c + \varepsilon_{s,c}, Y_{s,c} = b_s M_{s,c} + c_s' X_c + \eta_{s,c},$$

where $X_c$ denotes the clip-wise semantic feature value. The subject-level indirect effect was defined as $ab_s = a_s b_s$.

Group-level indirect effects were summarized by bootstrapping across subjects (1,000 iterations; mean and 95% confidence intervals). Statistical significance was assessed using clip-level permutation testing (5,000 permutations of $X_c$), yielding permutation-based $p$ values and $z$ statistics. False discovery rate correction was applied across all rich-club hub × target sub-network pairs.

### Supplementary

#### Supplementary figures

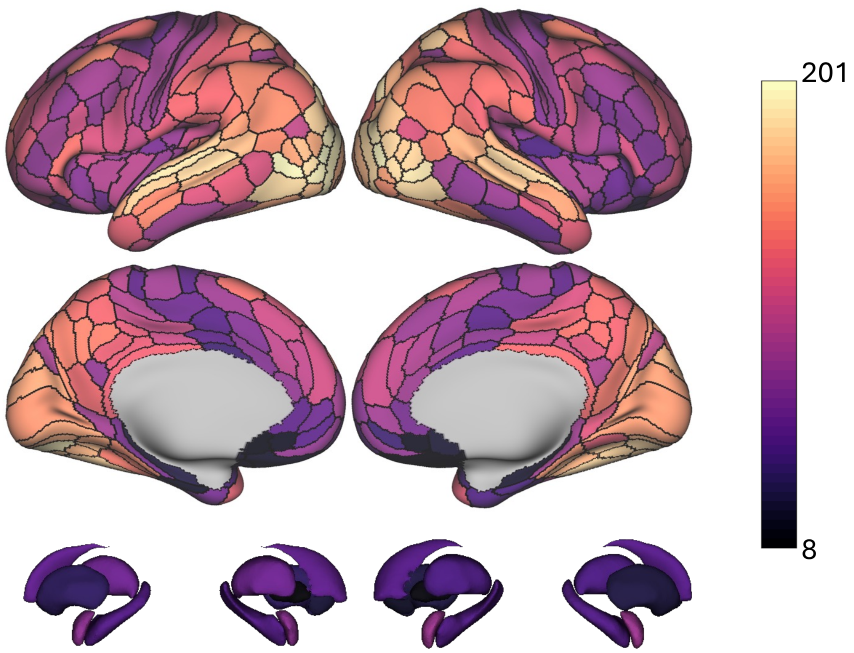


***Supplementary Fig. 1.* Node degree of the 374 ROIs, averaged from all clips retaining the top 30% strongest connections in the ISFC matrix.**

A. WordNet-based semantic labels across clips

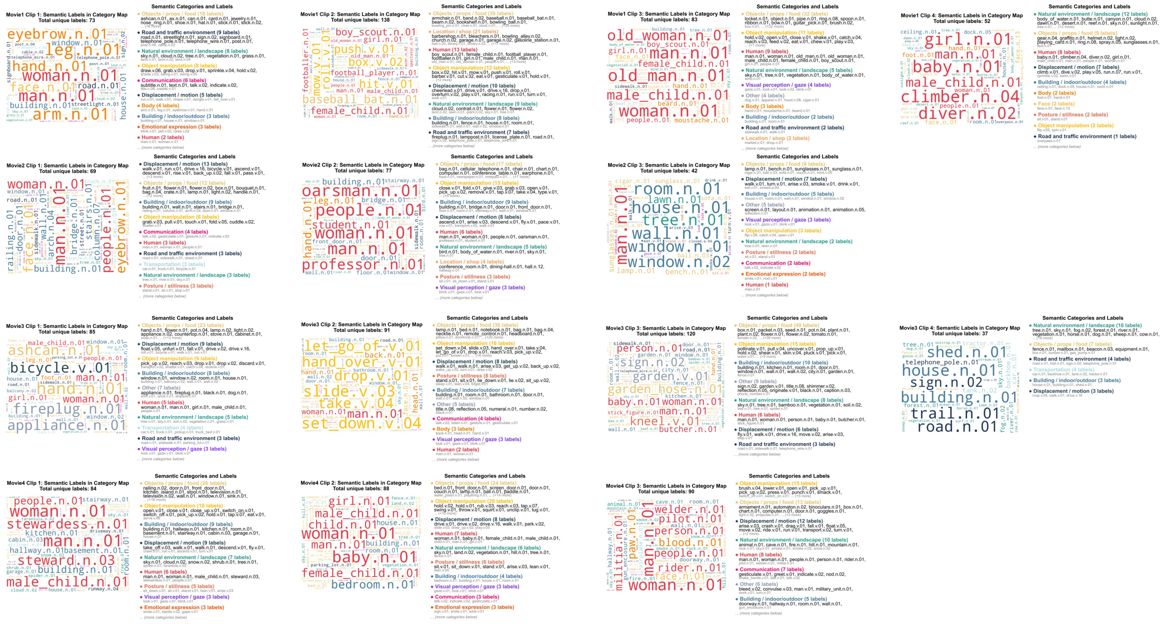

B. TR-wise human-related features across clips

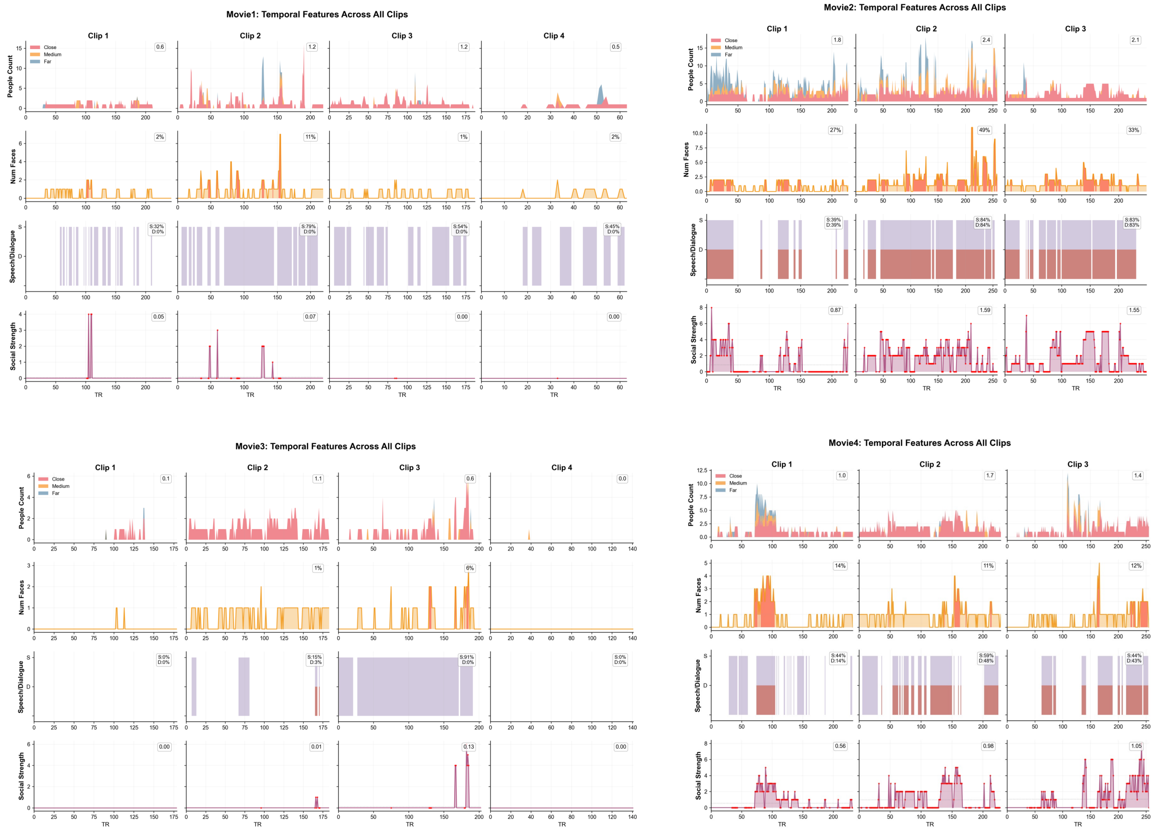

***Supplementary Fig. 2. Semantic and human-related stimulus features across movie clips.***

(A) WordNet-based semantic labels across clips. For each movie clip, TR-resolved semantic annotations from the HCP 7T movie-watching dataset were summarized as word clouds, illustrating the most frequently occurring WordNet label categories within each clip. (B) TR-wise human-related features across clips. For each clip, time-resolved traces show the presence and intensity of human-related features across TRs, including the number of detected people (stratified by viewing distance: close/medium/far), face counts, binary indicators of speech and dialogue, and a derived social-strength measure.

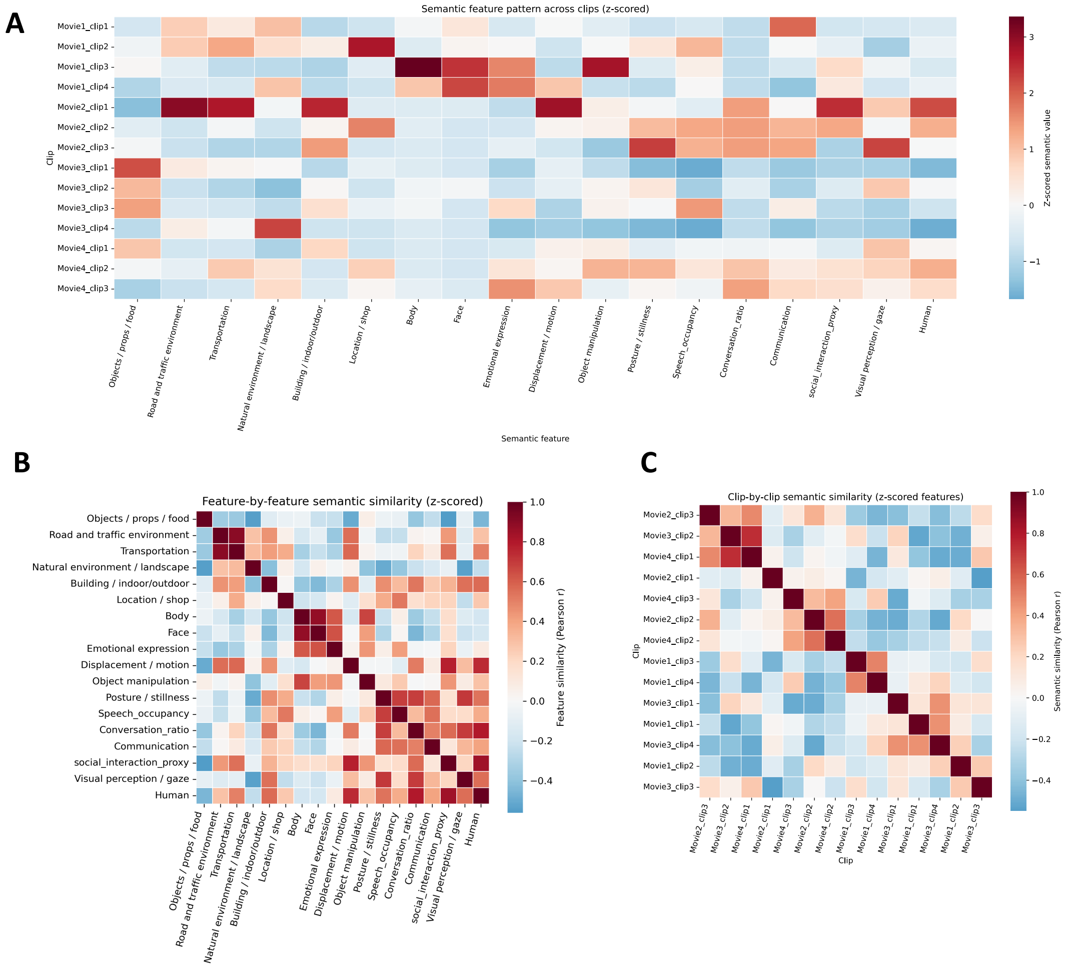


***Supplementary Fig. 3*. Stimulus structure of movie clips across runs.** (A) Clip-wise semantic profiles derived from TR-resolved annotations provided by the HCP 7T movie-watching dataset. Each movie clip was characterized along 18 predefined semantic dimensions. Values represent the mean feature strength across all TRs within each clip.Heatmap showing z-scored values of 18 predefined semantic dimensions for each movie. (B) Feature-by-feature semantic similarity. Pairwise Pearson correlation matrix between semantic features computed across clips using z-scored feature values. Positive correlations indicate semantic dimensions that tend to co-occur across clips (e.g., speech-related and social features), whereas negative correlations reflect opposing semantic profiles. The diagonal represents self-correlations. (C) Clip-by-clip semantic similarity. Pairwise Pearson correlation matrix between movie clips based on their full semantic feature profiles. Higher similarity values indicate clips with comparable semantic composition, revealing structured relationships both within and across movie runs.

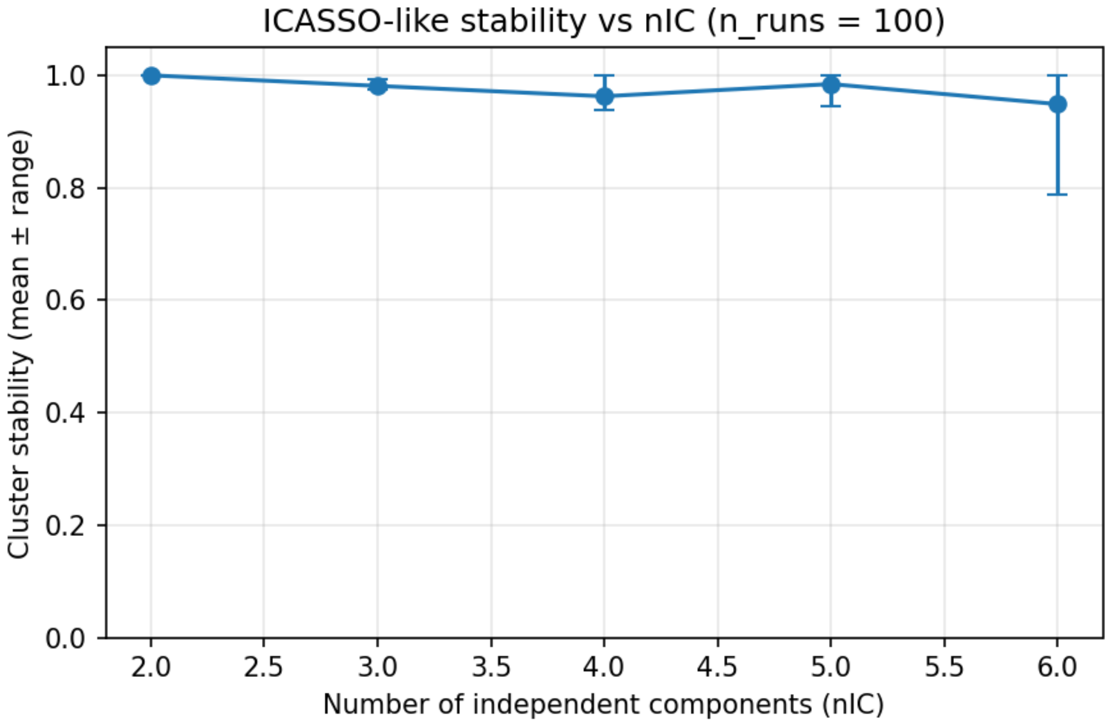


***Supplementary Fig. 4.*** **ICASSO-like stability analysis across different numbers of independent components ($n_{IC}$).**

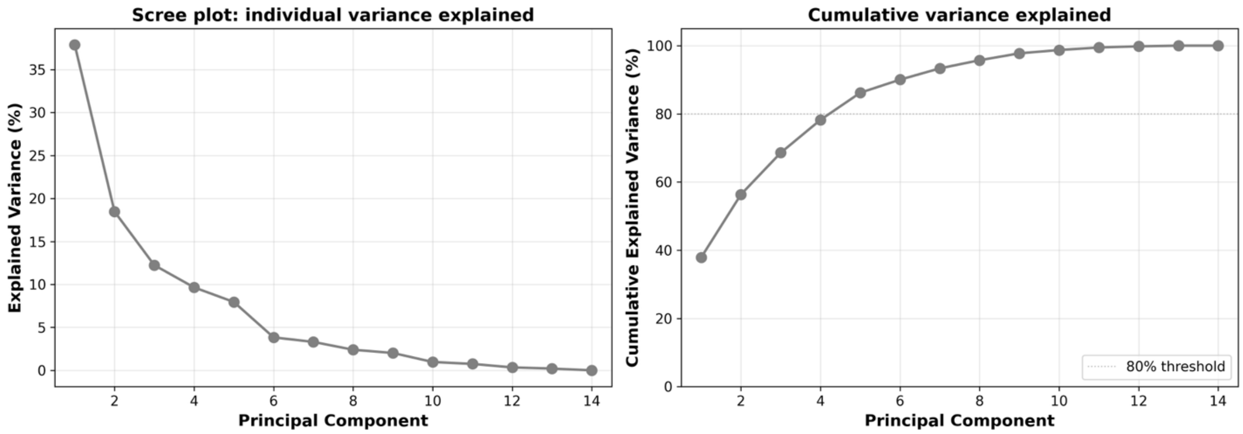


***Supplementary Fig. 5.*** **Variance explained by principal components of clip-wise stimulus Features**

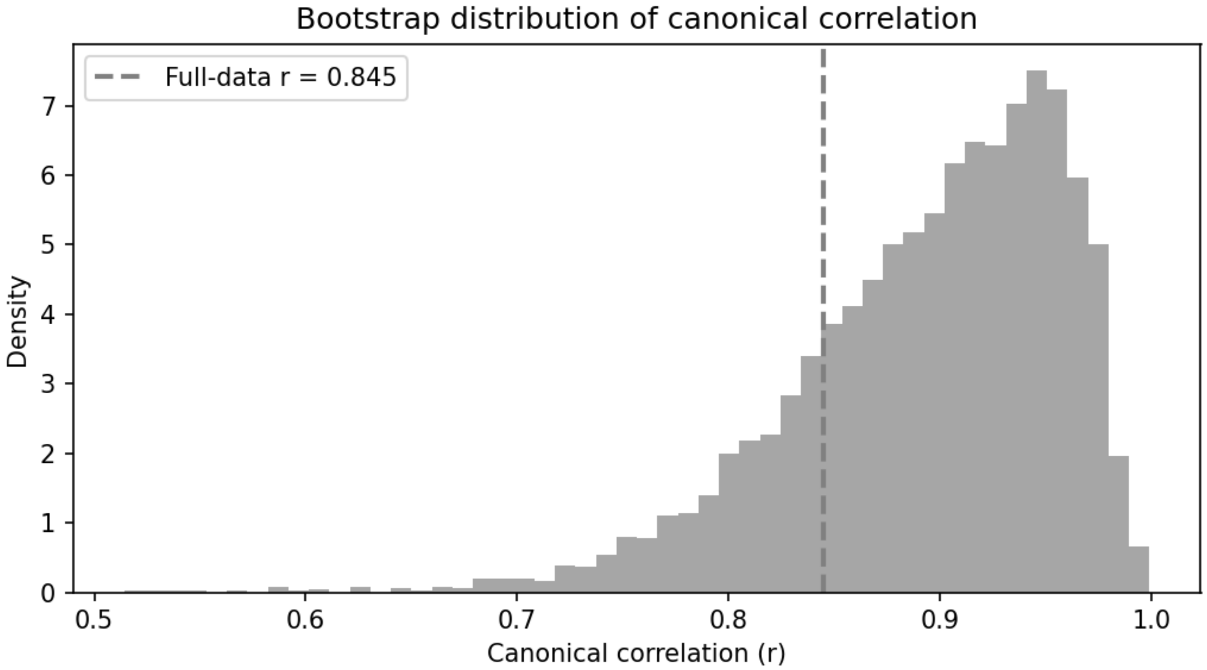


**Supplementary Fig. 6. Bootstrap distribution of canonical correlation from CCA.** Histogram showing the bootstrap distribution of the first canonical correlation coefficient obtained from canonical correlation analysis (CCA) between clip-wise brain and semantic feature representations. Bootstrap resampling was performed by repeatedly resampling clips with replacement and refitting the CCA model. The dashed vertical line indicates the canonical correlation estimated from the full dataset (r = 0.845). The distribution demonstrates the robustness and stability of the observed brain–stimulus coupling across resamples.

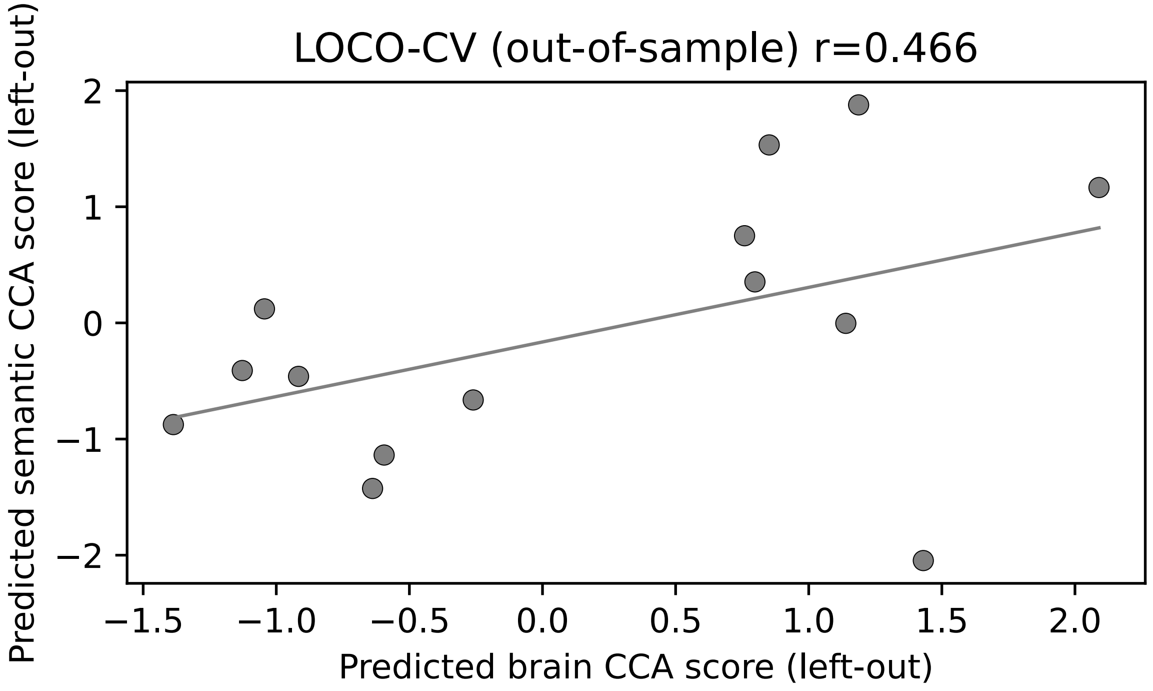


**Supplementary Fig. 7. LOCO-CV validation of the brain–stimulus canonical association.**

Leave-one-clip-out cross-validation (LOCO-CV) was used to assess the generalizability of the canonical correlation analysis (CCA) model. The positive correlation across clips (r=0.466) indicates that the brain–stimulus coupling captured by the CCA model generalizes to unseen stimuli.

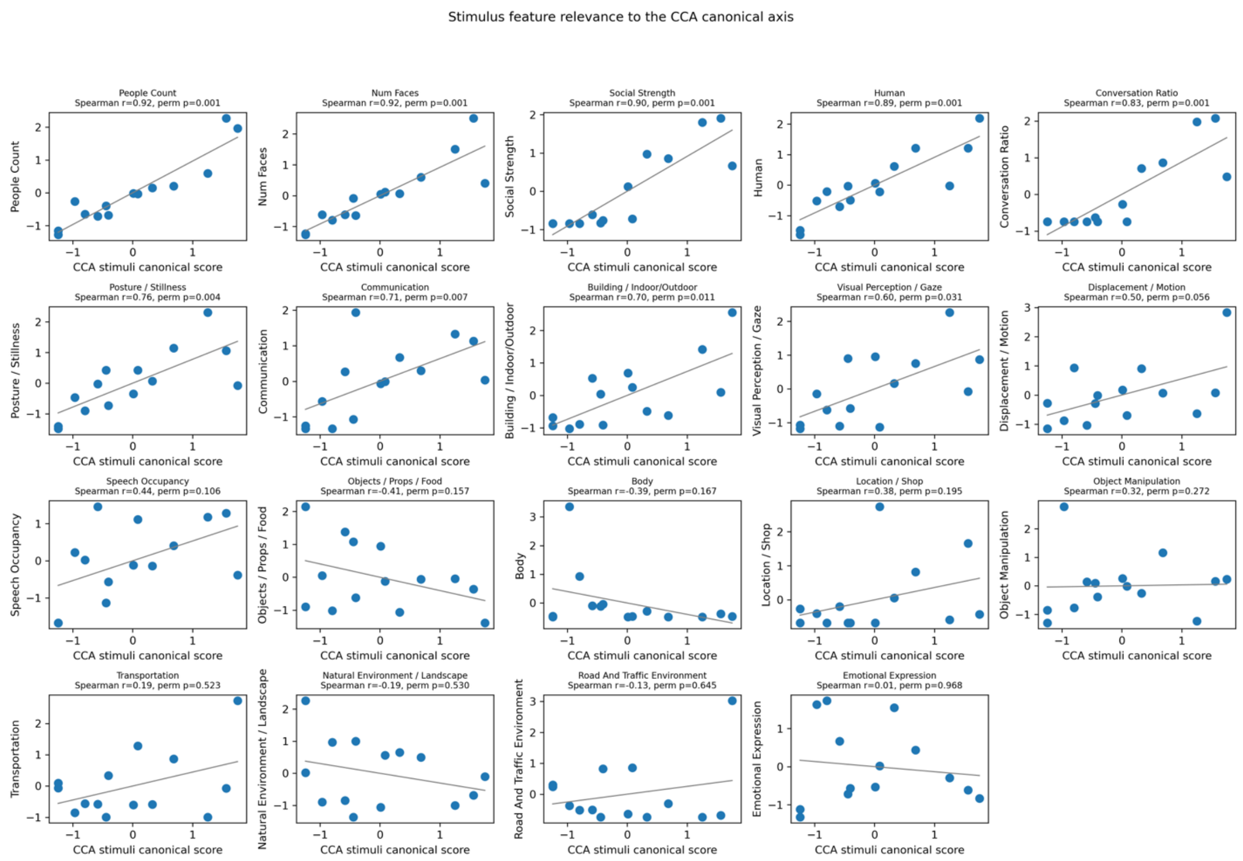


**Supplementary Fig. 8. Stimulus feature relevance to the CCA canonical axis.** Scatter plots showing the relationship between the CCA-derived semantic score and individual semantic feature measures across movie clips. Each panel corresponds to one stimuli feature. Each point represents one movie clip. The x-axis denotes the CCA semantic score obtained from canonical correlation analysis, and the y-axis denotes the z-scored value of the corresponding semantic feature. Solid lines indicate least-squares regression fits for visualization. Spearman rank correlation coefficients (r) and associated p values are reported in each panel, quantifying the monotonic relationship between individual semantic features and the CCA stimuli dimension.

Movie1_clip 1

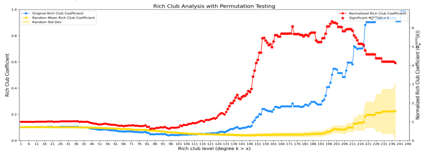


Movie3_clip1

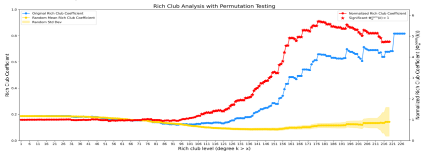


Movie1_clip 2

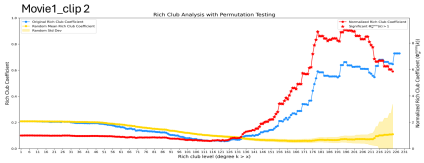


Movie3_clip2

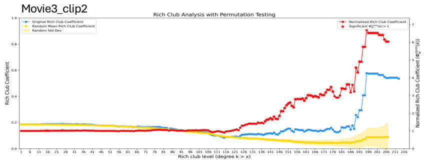


Movie1_clip 3

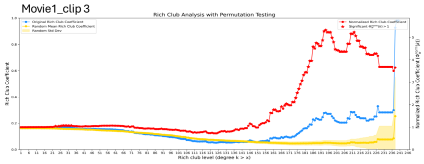


Movie3_clip3

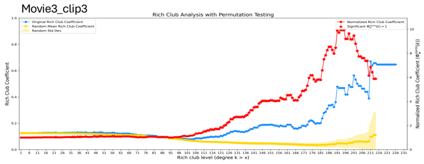


Movie1_clip 4

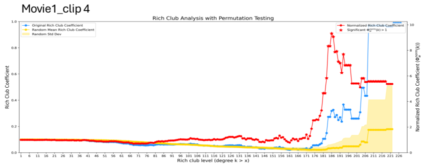


Movie3_clip4

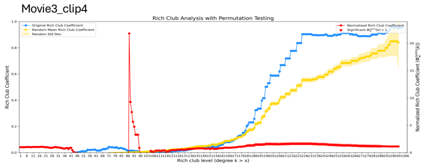


Movie 1_clip 5

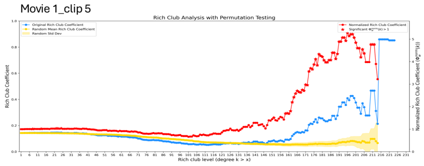


Movie3_clip5

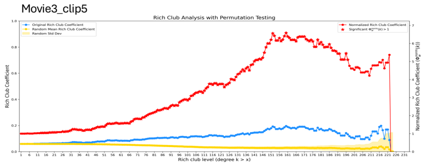


Movie 2_clip 1

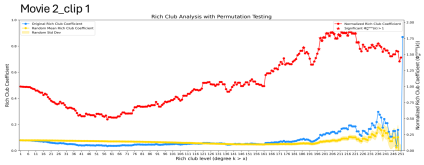


Movie4_clip1

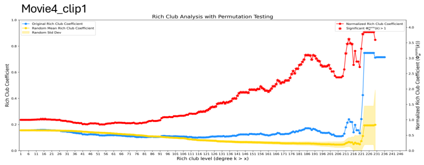


Movie 2_clip 2

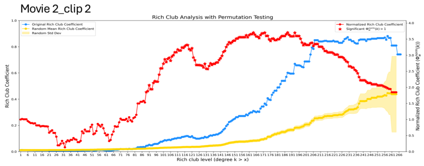


Movie4_clip2

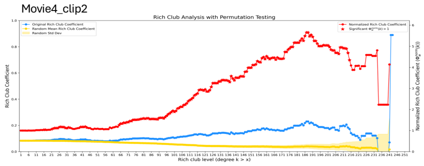


Movie 2_clip 3

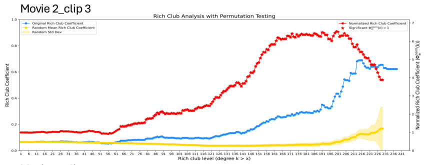


Movie4_clip3

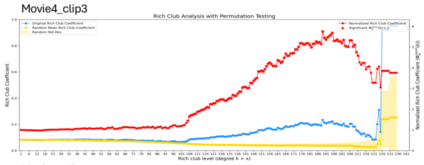


Movie 2_clip 4

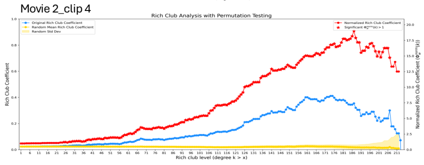


Movie4_clip4

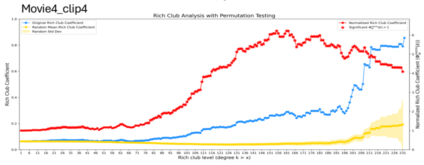

***Supplementary Fig. 9.*** *Rich-club analysis results of each clip based on group-averaged ISFC matrix. Each subplot shows the rich-club coefficient values for a range of k responses of the whole-brain network of each clip. Blue line: the original rich-club coefficient, yellow line: random network coefficient, and red line:normalized rich-club coefficient.*

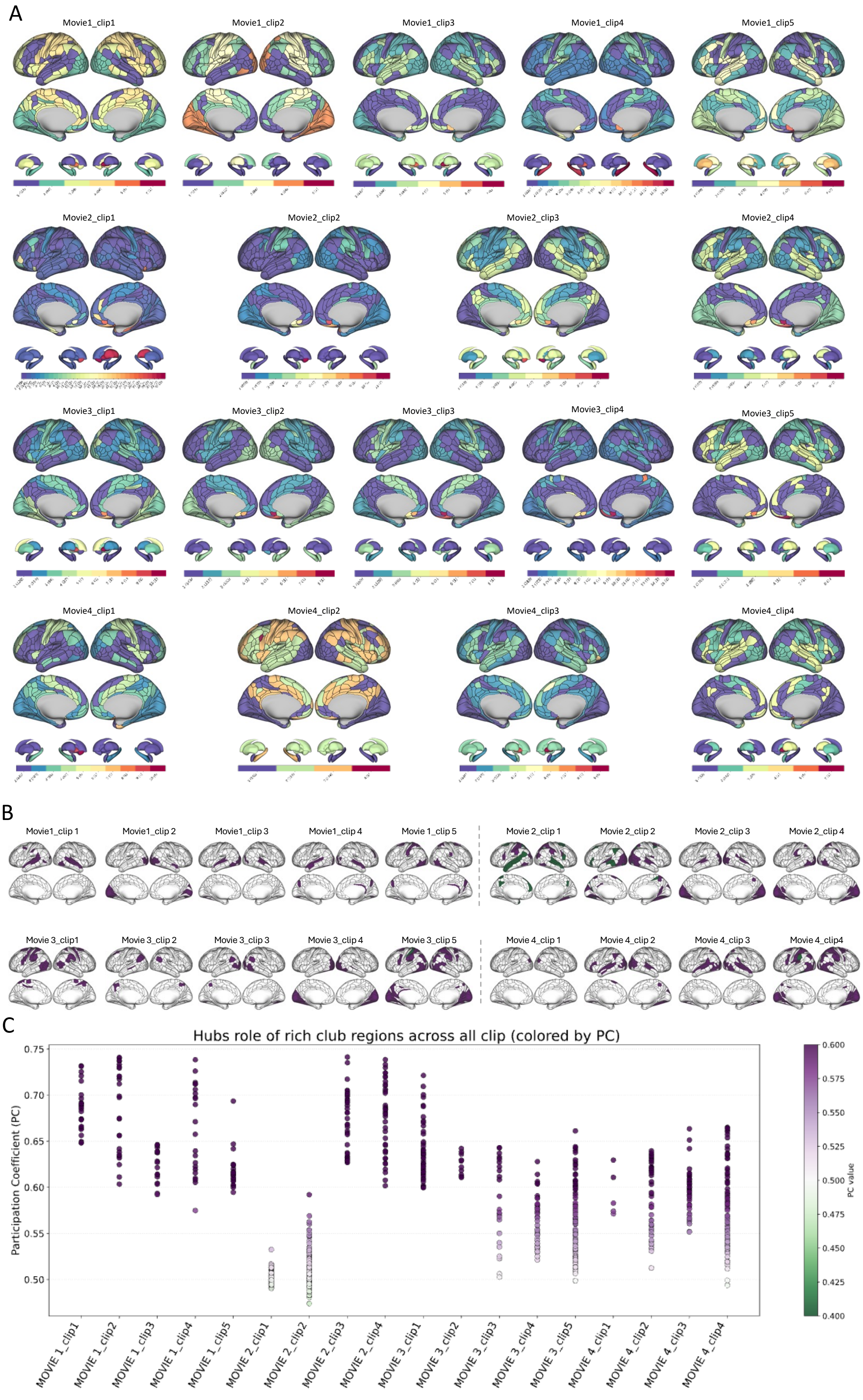
A
Movie1_clip1
Movie1_clip2
Movie1_clip3
Movie1_clip4
Movie1_clip5
Movie2_clip1
Movie2_clip2
Movie2_clip3
Movie2_clip4
Movie3_clip1
Movie3_clip2
Movie3_clip3
Movie3_clip4
Movie3_clip5
Movie4_clip1
Movie4_clip2
Movie4_clip3
Movie4_clip4
B
Movie1_clip 1
Movie1_clip 2
Movie1_clip 3
Movie1_clip 4
Movie 1_clip 5
Movie 2_clip 1
Movie 2_clip 2
Movie 2_clip 3
Movie 2_clip 4
Movie 3_clip1
Movie 3_clip 2
Movie 3_clip 3
Movie 3_clip 4
Movie 3_clip 5
Movie 4_clip 1
Movie 4_clip 2
Movie 4_clip 3
Movie 4_clip4
C
Hubs role of rich club regions across all clip (colored by PC)
Participation Coefficient (PC)
0.75
0.70
0.65
0.60
0.55
0.50
MOVIE 1_clip1
MOVIE 1_clip2
MOVIE 1_clip3
MOVIE 1_clip4
MOVIE 1_clip5
MOVIE 2_clip1
MOVIE 2_clip2
MOVIE 2_clip3
MOVIE 2_clip4
MOVIE 3_clip1
MOVIE 3_clip2
MOVIE 3_clip3
MOVIE 3_clip4
MOVIE 3_clip5
MOVIE 4_clip1
MOVIE 4_clip2
MOVIE 4_clip3
MOVIE 4_clip4
PC value
0.600
0.575
0.550
0.525
0.500
0.475
0.450
0.425
0.400

***Supplementary Fig. 10. Connter role of rich club regions.*** *(A) Community organization of each movie clip based on the Louvain community detection algorithm. For each clip, brain regions with the same color belong to the same community. The numbers shown in parentheses on the color bar represent the number of brain regions comprising the corresponding community. (B) participation coefficient map of rich-club regions for each clip. Purple: connector hubs ($PC > 0.5$); green: local hubs ($PC \leq 0.5$). (C) Distribution of PC values across clips.*

A

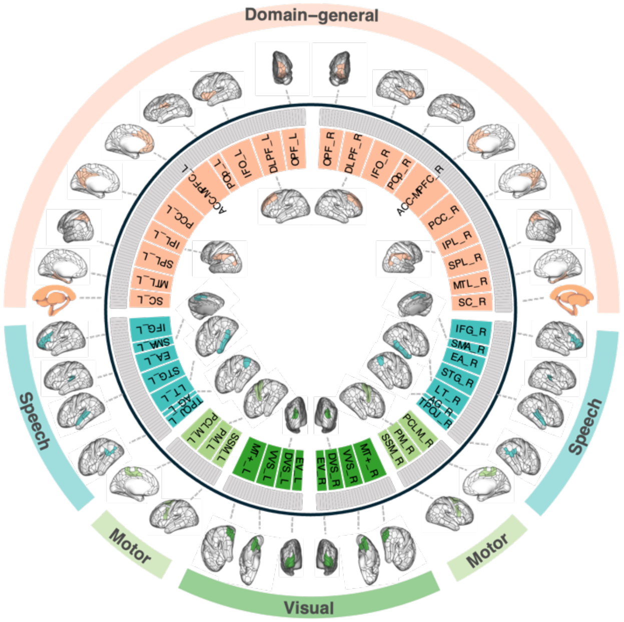
Domain–general
Speech
Speech
Motor
Motor
Visual
OPF_L
DLPF_L
IFO_L
POp_L
ACC-MPFC_L
PCC_L
IPL_L
SPL_L
MTL_L
SC_L
OPF_R
DLPF_R
IFO_R
POp_R
ACC-MPFC_R
PCC_R
IPL_R
SPL_R
MTL_R
SC_R
IFG_L
SMA_L
EA_L
STG_L
LT_L
AG_L
TPOJ_L
IFG_R
SMA_R
EA_R
STG_R
LT_R
AG_R
TPOJ_R
PCM_L
PM_L
SSM_L
PCM_R
PM_R
SSM_R
MT+_L
VVS_L
DVS_L
EV_L
MT+_R
VVS_R
DVS_R
EV_R

B

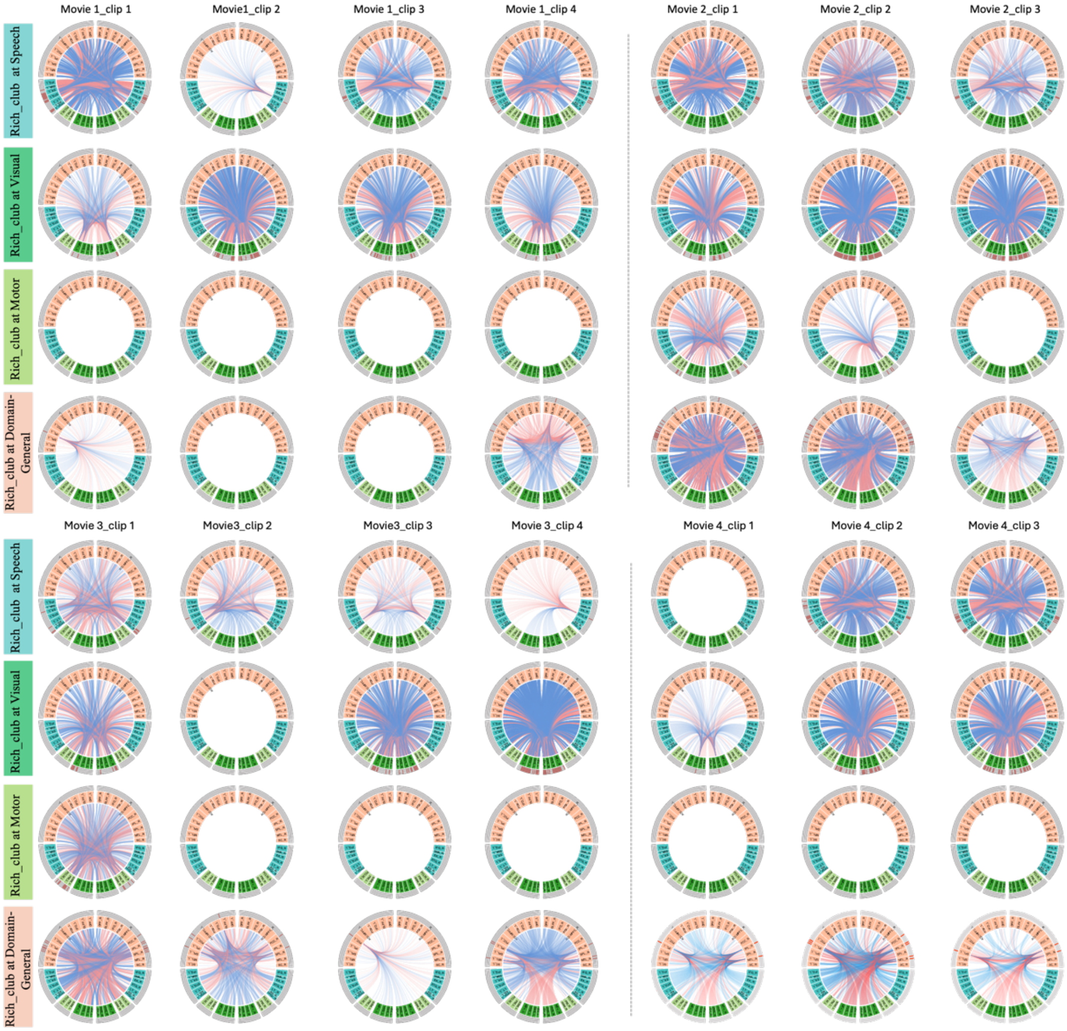
Movie 1_clip 1
Movie1_clip 2
Movie 1_clip 3
Movie 1_clip 4
Movie 2_clip 1
Movie 2_clip 2
Movie 2_clip 3
Rich_club at Speech
Rich_club at Visual
Rich_club at Motor
Rich_club at Domain-General
Movie 3_clip 1
Movie3_clip 2
Movie3_clip 3
Movie 3_clip 4
Movie 4_clip 1
Movie 4_clip 2
Movie 4_clip 3
Rich_club at Speech
Rich_club at Visual
Rich_club at Motor
Rich_club at Domain-General

272

***Supplementary Fig 11. Whole-brain connections of each clip's rich-club regions.*** *Rich-club regions are categorized into speech, visual, motor and domain-general systems based on their spatial locations and previously reported primary functions. (A) Categorization and anatomical locations of brain regions. (B) Connections between the rich-club regions and other non-rich-club regions are depicted using circular plots. Red: positive connections; blue: negative connections.*

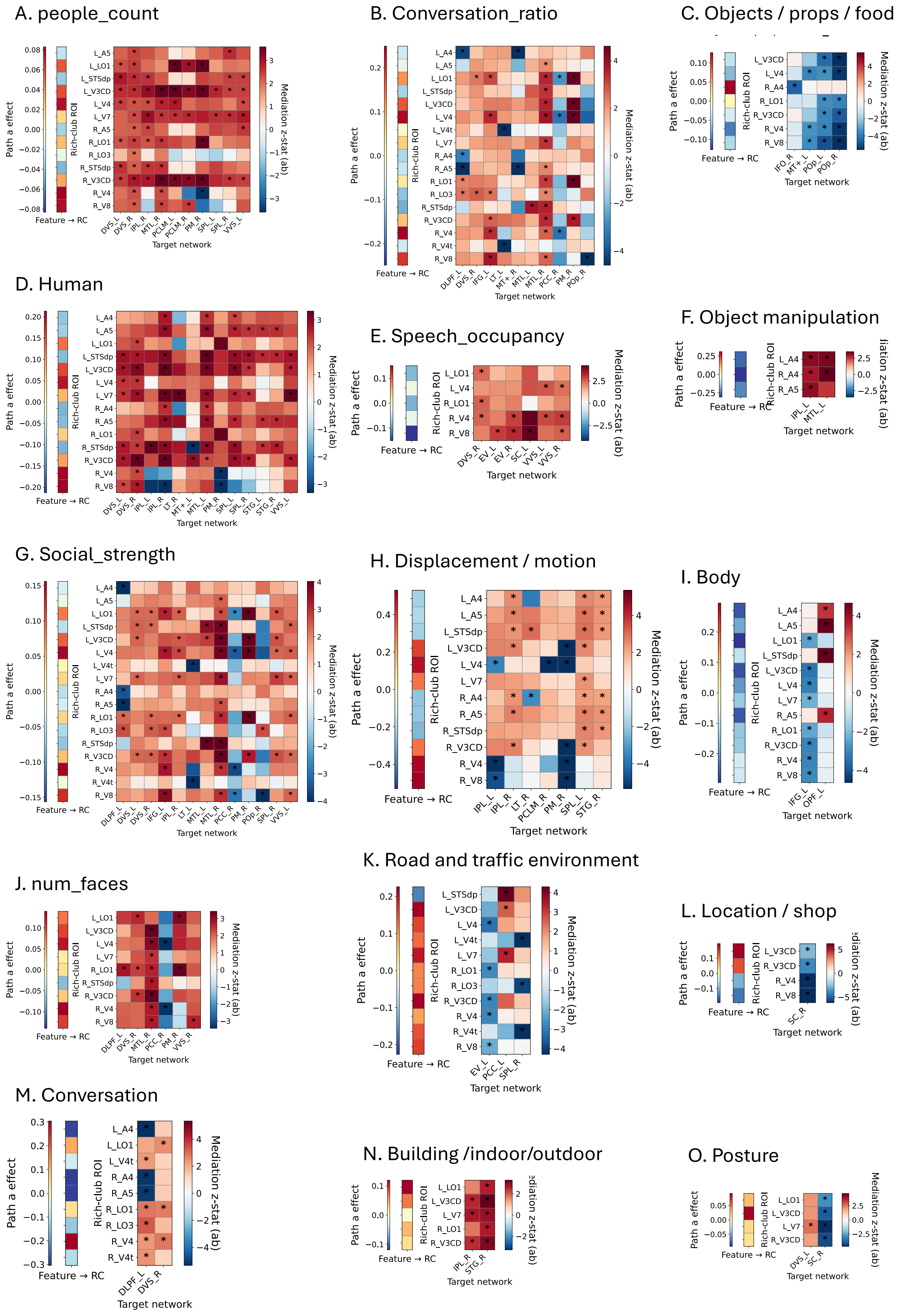
A. people_count
B. Conversation_ratio
C. Objects / props / food
D. Human
E. Speech_occupancy
F. Object manipulation
G. Social_strength
H. Displacement / motion
I. Body
J. num_faces
K. Road and traffic environment
L. Location / shop
M. Conversation
N. Building /indoor/outdoor
O. Posture
Path a effect
Rich-club ROI
Feature → RC
Target network
Mediation z-stat (ab)

***Supplementary Fig 12. Feature-specific effects on rich-club–mediated network integration during naturalistic movie viewing.*** *Heatmaps illustrate the path effects linking stimulus features to large-scale brain network integration through rich-club hubs. Each panel (A–O) corresponds to a stimulus feature extracted from the movie clips that exhibited significant mediation effects. The vertical bar on the left of each panel shows the path effect from the stimulus feature to the rich-club hubs (Feature → RC), while the main heatmap shows the mediation effects from rich-club hubs to target cortical networks (RC → network). Color values represent standardized mediation statistics (Z-statistics), with warm colors indicating positive effects and cool colors indicating negative effects. Rows correspond to rich-club hub regions, and columns correspond to target functional networks. * indicate statistically significant effects after correction for multiple comparisons ($p < 0.05$ ; FDR-corrected).*

### *Supplementary table*

***Supplementary Table 1***

***Description of Movie runs. Movie1 and Movie3 consist of clips from independent films available under a Creative Commons license, while Movie2 and Movie4 include clips from Hollywood films.***

| ***Clip number*** | ***Movie1*** | | ***Movie2*** | | ***Movie3*** | | ***Movie4*** | |
|---|---|---|---|---|---|---|---|---|
| | ***Leng (TRs)*** | ***Stimuli*** | ***Leng (TRs)*** | ***Stimuli*** | ***Leng (TRs)*** | ***Stimuli*** | ***Leng (TRs)*** | ***Stimuli*** |
| | ***20*** | *REST* | ***20*** | *REST* | ***20*** | *REST* | ***20*** | *REST* |
| ***Clip1*** | *244* | *Two Men (2009)* | *227* | *Inception (2010)* | *180* | *Off the Shelf: flower (2008)* | *233* | *Home Alone (1990)* |
| | ***20*** | *REST* | ***20*** | *REST* | ***20*** | *REST* | ***20*** | *REST* |
| ***Clip2*** | *220* | *Welcome to Bridgeville (2011)* | *259* | *The Social Network (2010)* | *184* | *1212: hotel (year unavailable)* | *230* | *Erin Brockovich (2000)* |
| | ***20*** | *REST* | ***20*** | *REST* | ***20*** | *REST* | ***20*** | *REST* |
| ***Clip3*** | *188* | *Pockets (2008)* | *250* | *Ocean's Eleven (2001)* | *204* | *Mrs. Meyer's Clean Day: garden (2013)* | *156* | *The Empire Strikes Back (1980)* |

| | | | | | | | | |
|---|---|---|---|---|---|---|---|---|
| | ***20*** | *REST* | ***20*** | *REST* | ***20*** | *REST* | ***20*** | *REST* |
| ***Clip4*** | *64* | *Inside the Human Body: overcome (2011)* | ***80 TRs*** | *Validation clip : Degrees South (2011); LXIV (2011)* | *142* | ***Clip4 :*** *Northwest Passage dreary: (year unavailable)* | ***80*** | *Validation clip: Degrees South (2011); LXIV (2011)* |
| | ***20*** | *REST* | ***20*** | *REST* | ***20*** | *REST* | ***20*** | *REST* |
| ***Clip5*** | ***80*** | *Validation clip : Degrees South (2011); LXIV (2011)* | | | ***80*** | *Validation clip: Degrees South (2011); LXIV (2011)* | | |
| | ***20*** | *REST* | | | ***20*** | *REST* | | |

***Supplementary Table 2***

***Global efficiency differences of ISFC between clip pairs within each movie, across network thresholds***

**Movie 1**

| Threshold | clip_1_vs_clip_2 | clip_1_vs_clip_3 | clip_1_vs_clip_4 | clip_1_vs_clip_5 | clip_2_vs_clip_3 | clip_2_vs_clip_4 | clip_2_vs_clip_5 | clip_3_vs_clip_4 | clip_3_vs_clip_5 | clip_4_vs_clip_5 |
|---|---|---|---|---|---|---|---|---|---|---|
| **10** | 0.022 | 0.035 | 0.000 | 0.002 | 0.013 | 0.023 | 0.025 | 0.035 | 0.037 | 0.002 |
| **20** | 0.023 | 0.026 | 0.012 | 0.013 | 0.003 | 0.035 | 0.036 | 0.038 | 0.039 | 0.001 |
| **30** | 0.025 | 0.026 | 0.010 | 0.014 | 0.001 | 0.036 | 0.040 | 0.037 | 0.040 | 0.004 |
| **40** | 0.026 | 0.026 | 0.012 | 0.015 | 0.000 | 0.038 | 0.041 | 0.038 | 0.041 | 0.003 |
| **50** | 0.026 | 0.026 | 0.013 | 0.015 | 0.001 | 0.039 | 0.041 | 0.039 | 0.041 | 0.002 |
| **60** | 0.026 | 0.026 | 0.014 | 0.015 | 0.001 | 0.040 | 0.041 | 0.039 | 0.041 | 0.002 |
| **70** | 0.026 | 0.026 | 0.014 | 0.015 | 0.001 | 0.040 | 0.041 | 0.039 | 0.041 | 0.002 |
| **80** | 0.026 | 0.026 | 0.014 | 0.015 | 0.001 | 0.040 | 0.041 | 0.039 | 0.041 | 0.002 |
| **90** | 0.026 | 0.026 | 0.014 | 0.015 | 0.001 | 0.040 | 0.041 | 0.039 | 0.041 | 0.002 |
| **100** | 0.026 | 0.026 | 0.014 | 0.015 | 0.001 | 0.040 | 0.041 | 0.039 | 0.041 | 0.002 |

**Movie 2**

| Threshold ( th% of network density) | clip_1_vs_clip_2 | clip_1_vs_clip_3 | clip_1_vs_clip_4 | clip_2_vs_clip_3 | clip_2_vs_clip_4 | clip_3_vs_clip_4 |
|---|---|---|---|---|---|---|
| **10** | 0.004 | 0.004 | 0.028 | 0.007 | 0.024 | 0.032 |
| **20** | 0.022 | 0.036 | 0.020 | 0.014 | 0.002 | 0.016 |
| **30** | 0.041 | 0.058 | 0.044 | 0.017 | 0.003 | 0.014 |
| **40** | 0.050 | 0.068 | 0.054 | 0.018 | 0.005 | 0.013 |
| **50** | 0.054 | 0.072 | 0.058 | 0.018 | 0.004 | 0.014 |
| **60** | 0.054 | 0.073 | 0.059 | 0.019 | 0.005 | 0.014 |
| **70** | 0.054 | 0.073 | 0.059 | 0.019 | 0.005 | 0.014 |
| **80** | 0.054 | 0.073 | 0.059 | 0.019 | 0.005 | 0.014 |
| **90** | 0.054 | 0.073 | 0.059 | 0.019 | 0.005 | 0.014 |
| **100** | 0.054 | 0.073 | 0.059 | 0.019 | 0.005 | 0.014 |

**Movie 3**

| Threshold ( th% of network density) | clip_1_vs_clip_2 | clip_1_vs_clip_3 | clip_1_vs_clip_4 | clip_1_vs_clip_5 | clip_2_vs_clip_3 | clip_2_vs_clip_4 | clip_2_vs_clip_5 | clip_3_vs_clip_4 | clip_3_vs_clip_5 | clip_4_vs_clip_5 |
|---|---|---|---|---|---|---|---|---|---|---|
| **10** | 0.021 | 0.011 | 0.046 | 0.012 | 0.010 | 0.067 | 0.008 | 0.057 | 0.002 | 0.058 |
| **20** | 0.025 | 0.017 | 0.049 | 0.019 | 0.008 | 0.074 | 0.006 | 0.066 | 0.001 | 0.067 |
| **30** | 0.027 | 0.018 | 0.046 | 0.019 | 0.009 | 0.074 | 0.009 | 0.065 | 0.000 | 0.065 |
| **40** | 0.028 | 0.018 | 0.047 | 0.018 | 0.010 | 0.075 | 0.010 | 0.065 | 0.000 | 0.065 |
| **50** | 0.027 | 0.018 | 0.048 | 0.018 | 0.010 | 0.075 | 0.010 | 0.065 | 0.000 | 0.065 |
| **60** | 0.027 | 0.018 | 0.048 | 0.018 | 0.009 | 0.075 | 0.009 | 0.066 | 0.000 | 0.066 |
| **70** | 0.028 | 0.018 | 0.048 | 0.018 | 0.010 | 0.075 | 0.010 | 0.066 | 0.000 | 0.066 |
| **80** | 0.028 | 0.018 | 0.048 | 0.018 | 0.010 | 0.075 | 0.010 | 0.066 | 0.000 | 0.066 |
| **90** | 0.028 | 0.018 | 0.048 | 0.018 | 0.010 | 0.075 | 0.010 | 0.066 | 0.000 | 0.066 |
| **100** | 0.028 | 0.018 | 0.048 | 0.018 | 0.010 | 0.075 | 0.010 | 0.066 | 0.000 | 0.066 |

**Movie 4**

| Threshold ( th% of network density) | clip_1_vs_clip_2 | clip_1_vs_clip_3 | clip_1_vs_clip_4 | clip_2_vs_clip_3 | clip_2_vs_clip_4 | clip_3_vs_clip_4 |
|---|---|---|---|---|---|---|
| **10** | 0.002 | 0.002 | 0.008 | 0.004 | 0.006 | 0.010 |
| **20** | 0.001 | 0.014 | 0.014 | 0.013 | 0.012 | 0.000 |
| **30** | 0.003 | 0.008 | 0.017 | 0.005 | 0.014 | 0.009 |
| **40** | 0.006 | 0.010 | 0.020 | 0.004 | 0.014 | 0.010 |
| **50** | 0.006 | 0.009 | 0.020 | 0.004 | 0.014 | 0.010 |
| **60** | 0.006 | 0.009 | 0.020 | 0.004 | 0.014 | 0.010 |
| **70** | 0.006 | 0.009 | 0.020 | 0.003 | 0.014 | 0.011 |
| **80** | 0.006 | 0.009 | 0.020 | 0.003 | 0.014 | 0.011 |
| **90** | 0.006 | 0.009 | 0.020 | 0.003 | 0.014 | 0.011 |
| **100** | 0.006 | 0.009 | 0.020 | 0.003 | 0.014 | 0.011 |

***Supplementary Table 3***

***Modularity Q scores between clip pairs within each movie, across network thresholds***

**Movie 1**

| Threshold | clip_1_vs_clip_2 | clip_1_vs_clip_3 | clip_1_vs_clip_4 | clip_1_vs_clip_5 | clip_2_vs_clip_3 | clip_2_vs_clip_4 | clip_2_vs_clip_5 | clip_3_vs_clip_4 | clip_3_vs_clip_5 | clip_4_vs_clip_5 |
|---|---|---|---|---|---|---|---|---|---|---|
| **10** | 0.009 | 0.000 | 0.016 | 0.041 | 0.009 | 0.025 | 0.032 | 0.015 | 0.041 | 0.057 |
| **20** | 0.020 | 0.045 | 0.029 | 0.013 | 0.065 | 0.049 | 0.007 | 0.016 | 0.058 | 0.042 |
| **30** | 0.008 | 0.070 | 0.028 | 0.005 | 0.078 | 0.036 | 0.013 | 0.042 | 0.065 | 0.022 |
| **40** | 0.005 | 0.083 | 0.025 | 0.015 | 0.088 | 0.030 | 0.020 | 0.058 | 0.068 | 0.010 |
| **50** | 0.013 | 0.083 | 0.031 | 0.022 | 0.096 | 0.044 | 0.035 | 0.052 | 0.060 | 0.009 |
| **60** | 0.023 | 0.068 | 0.020 | 0.006 | 0.091 | 0.043 | 0.029 | 0.048 | 0.062 | 0.014 |
| **70** | 0.016 | 0.076 | 0.036 | 0.013 | 0.092 | 0.053 | 0.030 | 0.039 | 0.062 | 0.023 |
| **80** | 0.032 | 0.072 | 0.033 | 0.010 | 0.104 | 0.065 | 0.042 | 0.040 | 0.062 | 0.022 |
| **90** | 0.027 | 0.075 | 0.035 | 0.005 | 0.102 | 0.062 | 0.032 | 0.039 | 0.070 | 0.031 |
| **100** | 0.002 | 0.104 | 0.064 | 0.042 | 0.102 | 0.062 | 0.039 | 0.039 | 0.062 | 0.023 |

**Movie 2**

| Threshold ( th% of network density) | clip_1_vs_clip_2 | clip_1_vs_clip_3 | clip_1_vs_clip_4 | clip_2_vs_clip_3 | clip_2_vs_clip_4 | clip_3_vs_clip_4 |
|---|---|---|---|---|---|---|
| **10** | 0.096 | 0.007 | 0.000 | 0.088 | 0.096 | 0.007 |
| **20** | 0.040 | 0.060 | 0.059 | 0.020 | 0.018 | 0.002 |
| **30** | 0.019 | 0.074 | 0.076 | 0.055 | 0.057 | 0.002 |
| **40** | 0.010 | 0.082 | 0.099 | 0.072 | 0.088 | 0.016 |

| | | | | | | |
|---|---|---|---|---|---|---|
| **50** | 0.014 | 0.113 | 0.114 | 0.099 | 0.100 | 0.001 |
| **60** | 0.025 | 0.112 | 0.116 | 0.086 | 0.091 | 0.005 |
| **70** | 0.028 | 0.125 | 0.107 | 0.097 | 0.079 | 0.018 |
| **80** | 0.023 | 0.108 | 0.102 | 0.085 | 0.079 | 0.006 |
| **90** | 0.023 | 0.111 | 0.103 | 0.088 | 0.080 | 0.008 |
| **100** | 0.027 | 0.111 | 0.102 | 0.084 | 0.075 | 0.008 |

**Movie 3**

| Threshold ( th% of network density) | clip_1_vs_clip_2 | clip_1_vs_clip_3 | clip_1_vs_clip_4 | clip_1_vs_clip_5 | clip_2_vs_clip_3 | clip_2_vs_clip_4 | clip_2_vs_clip_5 | clip_3_vs_clip_4 | clip_3_vs_clip_5 | clip_4_vs_clip_5 |
|---|---|---|---|---|---|---|---|---|---|---|
| **10** | 0.077 | 0.063 | 0.025 | 0.077 | 0.014 | 0.052 | 0.000 | 0.038 | 0.014 | 0.052 |
| **20** | 0.080 | 0.069 | 0.143 | 0.081 | 0.011 | 0.063 | 0.001 | 0.074 | 0.011 | 0.062 |
| **30** | 0.080 | 0.074 | 0.143 | 0.085 | 0.006 | 0.063 | 0.005 | 0.069 | 0.011 | 0.059 |
| **40** | 0.094 | 0.053 | 0.154 | 0.071 | 0.041 | 0.060 | 0.023 | 0.101 | 0.018 | 0.083 |
| **50** | 0.105 | 0.077 | 0.198 | 0.093 | 0.029 | 0.093 | 0.013 | 0.122 | 0.016 | 0.106 |
| **60** | 0.096 | 0.050 | 0.196 | 0.064 | 0.046 | 0.101 | 0.032 | 0.146 | 0.014 | 0.132 |
| **70** | 0.083 | 0.041 | 0.202 | 0.083 | 0.042 | 0.119 | 0.000 | 0.160 | 0.041 | 0.119 |
| **80** | 0.065 | 0.042 | 0.202 | 0.068 | 0.024 | 0.137 | 0.003 | 0.160 | 0.027 | 0.134 |
| **90** | 0.061 | 0.048 | 0.199 | 0.065 | 0.013 | 0.138 | 0.004 | 0.151 | 0.017 | 0.134 |
| **100** | 0.071 | 0.032 | 0.193 | 0.074 | 0.038 | 0.122 | 0.003 | 0.160 | 0.041 | 0.119 |

**Movie 4**

| Threshold ( th% of network density) | clip_1_vs_clip_2 | clip_1_vs_clip_3 | clip_1_vs_clip_4 | clip_2_vs_clip_3 | clip_2_vs_clip_4 | clip_3_vs_clip_4 |
|---|---|---|---|---|---|---|
| **10** | 0.077 | 0.037 | 0.046 | 0.040 | 0.032 | 0.009 |
| **20** | 0.033 | 0.038 | 0.052 | 0.005 | 0.019 | 0.013 |
| **30** | 0.026 | 0.005 | 0.038 | 0.031 | 0.065 | 0.034 |
| **40** | 0.026 | 0.005 | 0.046 | 0.031 | 0.071 | 0.040 |
| **50** | 0.027 | 0.003 | 0.055 | 0.030 | 0.082 | 0.052 |
| **60** | 0.006 | 0.009 | 0.020 | 0.004 | 0.014 | 0.010 |
| **70** | 0.006 | 0.009 | 0.020 | 0.003 | 0.014 | 0.011 |
| **80** | 0.006 | 0.009 | 0.020 | 0.003 | 0.014 | 0.011 |
| **90** | 0.006 | 0.009 | 0.020 | 0.003 | 0.014 | 0.011 |
| **100** | 0.006 | 0.009 | 0.020 | 0.003 | 0.014 | 0.011 |

***Supplementary Table 4***

*Node-specific metrics of average degree and Coefficient variation*

| Lobe_name | ROI_label | Degree valua | Coefficient variation |
|---|---|---|---|
| Primary_Visual_L | L_V1 | 154 | 0.31 |
| MT+_Complex_and_Neighboring_Visual_Areas_L | L_MST | 160 | 0.27 |
| Dorsal_Stream_Visual_L | L_V6 | 92 | 0.41 |
| Early_Visual_L | L_V2 | 163 | 0.28 |
| Early_Visual_L | L_V3 | 165 | 0.35 |
| Early_Visual_L | L_V4 | 196 | 0.20 |
| Ventral_Stream_Visual_L | L_V8 | 176 | 0.29 |
| Somatosensory_and_Motor_L | L_4 | 86 | 0.52 |
| Somatosensory_and_Motor_L | L_3b | 82 | 0.52 |
| Premotor_L | L_FEF | 64 | 0.55 |
| Premotor_L | L_PEF | 85 | 0.62 |
| Premotor_L | L_55b | 113 | 0.43 |
| Dorsal_Stream_Visual_L | L_V3A | 146 | 0.32 |
| Posterior_Cingulate_L | L_RSC | 128 | 0.30 |
| Posterior_Cingulate_L | L_POS2 | 145 | 0.31 |
| Dorsal_Stream_Visual_L | L_V7 | 193 | 0.22 |
| Dorsal_Stream_Visual_L | L_IPS1 | 175 | 0.22 |
| Ventral_Stream_Visual_L | L_FFC | 162 | 0.33 |
| Dorsal_Stream_Visual_L | L_V3B | 169 | 0.33 |
| MT+_Complex_and_Neighboring_Visual_Areas_L | L_LO1 | 201 | 0.16 |
| MT+_Complex_and_Neighboring_Visual_Areas_L | L_LO2 | 196 | 0.21 |
| Ventral_Stream_Visual_L | L_PIT | 173 | 0.28 |
| MT+_Complex_and_Neighboring_Visual_Areas_L | L_MT | 172 | 0.26 |
| Early_Auditory_L | L_A1 | 136 | 0.30 |

| | | | |
|---|---|---|---|
| Temporo-Parieto-Occipital_Junction_L | L_PSL | 139 | 0.37 |
| Dorsolateral_Prefrontal_L | L_SFL | 137 | 0.32 |
| Posterior_Cingulate_L | L_PCV | 132 | 0.33 |
| Temporo-Parieto-Occipital_Junction_L | L_STV | 157 | 0.25 |
| Superior_Parietal_L | L_7Pm | 138 | 0.39 |
| Posterior_Cingulate_L | L_7m | 145 | 0.22 |
| Posterior_Cingulate_L | L_POS1 | 133 | 0.41 |
| Posterior_Cingulate_L | L_23d | 108 | 0.38 |
| Posterior_Cingulate_L | L_v23ab | 80 | 0.40 |
| Posterior_Cingulate_L | L_d23ab | 125 | 0.34 |
| Posterior_Cingulate_L | L_31pv | 127 | 0.26 |
| Paracentral_Lobular_and_Mid_Cingulate_L | L_5m | 56 | 0.54 |
| Paracentral_Lobular_and_Mid_Cingulate_L | L_5mv | 96 | 0.38 |
| Paracentral_Lobular_and_Mid_Cingulate_L | L_23c | 108 | 0.35 |
| Paracentral_Lobular_and_Mid_Cingulate_L | L_5L | 99 | 0.43 |
| Paracentral_Lobular_and_Mid_Cingulate_L | L_24dd | 62 | 0.51 |
| Paracentral_Lobular_and_Mid_Cingulate_L | L_24dv | 52 | 0.59 |
| Superior_Parietal_L | L_7AL | 148 | 0.33 |
| Paracentral_Lobular_and_Mid_Cingulate_L | L_SCEF | 86 | 0.41 |
| Paracentral_Lobular_and_Mid_Cingulate_L | L_6ma | 91 | 0.62 |
| Superior_Parietal_L | L_7Am | 131 | 0.35 |
| Superior_Parietal_L | L_7PL | 139 | 0.31 |
| Superior_Parietal_L | L_7PC | 149 | 0.36 |
| Superior_Parietal_L | L_LIPv | 155 | 0.27 |
| Superior_Parietal_L | L_VIP | 166 | 0.21 |
| Superior_Parietal_L | L_MIP | 127 | 0.32 |
| Somatosensory_and_Motor_L | L_1 | 91 | 0.44 |
| Somatosensory_and_Motor_L | L_2 | 126 | 0.50 |
| Somatosensory_and_Motor_L | L_3a | 89 | 0.38 |
| Premotor_L | L_6d | 62 | 0.54 |
| Paracentral_Lobular_and_Mid_Cingulate_L | L_6mp | 54 | 0.68 |
| Premotor_L | L_6v | 100 | 0.59 |
| Anterior_Cingulate_and_Medial_Prefrontal_L | L_p24pr | 55 | 0.51 |
| Anterior_Cingulate_and_Medial_Prefrontal_L | L_33pr | 41 | 1.01 |
| Anterior_Cingulate_and_Medial_Prefrontal_L | L_a24pr | 73 | 0.64 |
| Anterior_Cingulate_and_Medial_Prefrontal_L | L_p32pr | 72 | 0.59 |
| Anterior_Cingulate_and_Medial_Prefrontal_L | L_a24 | 56 | 0.58 |
| Anterior_Cingulate_and_Medial_Prefrontal_L | L_d32 | 103 | 0.33 |
| Anterior_Cingulate_and_Medial_Prefrontal_L | L_8BM | 109 | 0.39 |
| Anterior_Cingulate_and_Medial_Prefrontal_L | L_p32 | 62 | 0.54 |
| Anterior_Cingulate_and_Medial_Prefrontal_L | L_10r | 54 | 0.79 |
| Orbital_and_Polar_Frontal_L | L_47m | 50 | 0.61 |
| Dorsolateral_Prefrontal_L | L_8Av | 149 | 0.22 |
| Dorsolateral_Prefrontal_L | L_8Ad | 88 | 0.31 |
| Anterior_Cingulate_and_Medial_Prefrontal_L | L_9m | 112 | 0.33 |
| Dorsolateral_Prefrontal_L | L_8BL | 121 | 0.33 |
| Dorsolateral_Prefrontal_L | L_9p | 107 | 0.30 |
| Orbital_and_Polar_Frontal_L | L_10d | 89 | 0.40 |
| Dorsolateral_Prefrontal_L | L_8C | 88 | 0.51 |
| Inferior_Frontal_L | L_44 | 117 | 0.39 |
| Inferior_Frontal_L | L_45 | 138 | 0.35 |
| Inferior_Frontal_L | L_47l | 121 | 0.39 |
| Inferior_Frontal_L | L_a47r | 94 | 0.48 |
| Premotor_L | L_6r | 115 | 0.43 |
| Inferior_Frontal_L | L_IFJa | 81 | 0.54 |
| Inferior_Frontal_L | L_IFJp | 85 | 0.51 |
| Inferior_Frontal_L | L_IFSp | 112 | 0.41 |
| Inferior_Frontal_L | L_IFSa | 88 | 0.57 |
| Dorsolateral_Prefrontal_L | L_p9-46v | 80 | 0.48 |
| Dorsolateral_Prefrontal_L | L_46 | 111 | 0.45 |
| Dorsolateral_Prefrontal_L | L_a9-46v | 124 | 0.34 |
| Dorsolateral_Prefrontal_L | L_9-46d | 117 | 0.48 |
| Dorsolateral_Prefrontal_L | L_9a | 141 | 0.20 |
| Anterior_Cingulate_and_Medial_Prefrontal_L | L_10v | 81 | 0.45 |
| Orbital_and_Polar_Frontal_L | L_a10p | 107 | 0.36 |
| Orbital_and_Polar_Frontal_L | L_10pp | 61 | 0.55 |
| Orbital_and_Polar_Frontal_L | L_11l | 101 | 0.40 |
| Orbital_and_Polar_Frontal_L | L_13l | 51 | 0.66 |
| Orbital_and_Polar_Frontal_L | L_OFC | 19 | 0.97 |
| Orbital_and_Polar_Frontal_L | L_47s | 93 | 0.39 |
| Superior_Parietal_L | L_LIPd | 88 | 0.46 |
| Premotor_L | L_6a | 136 | 0.37 |
| Dorsolateral_Prefrontal_L | L_i6-8 | 89 | 0.43 |
| Dorsolateral_Prefrontal_L | L_s6-8 | 87 | 0.39 |
| Posterior_Opercular_L | L_43 | 108 | 0.37 |

| | | | |
|---|---|---|---|
| Posterior_Opercular_L | L_OP4 | 124 | 0.27 |
| Posterior_Opercular_L | L_OP1 | 99 | 0.41 |
| Posterior_Opercular_L | L_OP2-3 | 81 | 0.38 |
| Early_Auditory_L | L_52 | 85 | 0.34 |
| Early_Auditory_L | L_RI | 107 | 0.33 |
| Early_Auditory_L | L_PFcm | 131 | 0.38 |
| Insular_and_Frontal_Opercular_L | L_PoI2 | 101 | 0.42 |
| Auditory_Association_L | L_TA2 | 137 | 0.37 |
| Insular_and_Frontal_Opercular_L | L_FOP4 | 106 | 0.40 |
| Insular_and_Frontal_Opercular_L | L_MI | 96 | 0.45 |
| Insular_and_Frontal_Opercular_L | L_Pir | 47 | 0.61 |
| Insular_and_Frontal_Opercular_L | L_AVI | 74 | 0.60 |
| Insular_and_Frontal_Opercular_L | L_AAIC | 63 | 0.59 |
| Posterior_Opercular_L | L_FOP1 | 105 | 0.36 |
| Insular_and_Frontal_Opercular_L | L_FOP3 | 83 | 0.39 |
| Insular_and_Frontal_Opercular_L | L_FOP2 | 81 | 0.59 |
| Inferior_Parietal_L | L_PFt | 137 | 0.49 |
| Superior_Parietal_L | L_AIP | 132 | 0.42 |
| Medial_Temporal_L | L_EC | 18 | 1.01 |
| Medial_Temporal_L | L_PreS | 69 | 0.53 |
| Medial_Temporal_L | L_H | 29 | 0.89 |
| Posterior_Cingulate_L | L_ProS | 88 | 0.42 |
| Medial_Temporal_L | L_PeEc | 65 | 0.66 |
| Auditory_Association_L | L_STGa | 163 | 0.31 |
| Early_Auditory_L | L_PBelt | 150 | 0.30 |
| Auditory_Association_L | L_A5 | 190 | 0.24 |
| Medial_Temporal_L | L_PHA1 | 150 | 0.30 |
| Medial_Temporal_L | L_PHA3 | 142 | 0.26 |
| Auditory_Association_L | L_STSda | 178 | 0.23 |
| Auditory_Association_L | L_STSdp | 184 | 0.26 |
| Auditory_Association_L | L_STSvp | 169 | 0.27 |
| Lateral_Temporal_L | L_TGd | 128 | 0.36 |
| Lateral_Temporal_L | L_TE1a | 147 | 0.30 |
| Lateral_Temporal_L | L_TE1p | 124 | 0.31 |
| Lateral_Temporal_L | L_TE2a | 88 | 0.40 |
| Medial_Temporal_L | L_TF | 101 | 0.41 |
| Lateral_Temporal_L | L_TE2p | 99 | 0.50 |
| Lateral_Temporal_L | L_PHT | 117 | 0.36 |
| MT+_Complex_and_Neighboring_Visual_Areas_L | L_PH | 186 | 0.21 |
| Temporo-Parieto-Occipital_Junction_L | L_TPOJ1 | 172 | 0.26 |
| Temporo-Parieto-Occipital_Junction_L | L_TPOJ2 | 149 | 0.25 |
| Temporo-Parieto-Occipital_Junction_L | L_TPOJ3 | 106 | 0.45 |
| Posterior_Cingulate_L | L_DVT | 149 | 0.26 |
| Inferior_Parietal_L | L_PGp | 183 | 0.21 |
| Inferior_Parietal_L | L_IP2 | 125 | 0.41 |
| Inferior_Parietal_L | L_IP1 | 123 | 0.28 |
| Inferior_Parietal_L | L_IP0 | 176 | 0.28 |
| Inferior_Parietal_L | L_PFop | 133 | 0.41 |
| Inferior_Parietal_L | L_PF | 132 | 0.36 |
| Inferior_Parietal_L | L_PFm | 148 | 0.30 |
| Inferior_Parietal_L | L_PGi | 151 | 0.24 |
| Inferior_Parietal_L | L_PGs | 136 | 0.38 |
| Dorsal_Stream_Visual_L | L_V6A | 168 | 0.20 |
| Ventral_Stream_Visual_L | L_VMV1 | 161 | 0.33 |
| Ventral_Stream_Visual_L | L_VMV3 | 168 | 0.29 |
| Medial_Temporal_L | L_PHA2 | 144 | 0.26 |
| MT+_Complex_and_Neighboring_Visual_Areas_L | L_V4t | 186 | 0.27 |
| MT+_Complex_and_Neighboring_Visual_Areas_L | L_FST | 199 | 0.13 |
| MT+_Complex_and_Neighboring_Visual_Areas_L | L_V3CD | 196 | 0.25 |
| MT+_Complex_and_Neighboring_Visual_Areas_L | L_LO3 | 164 | 0.31 |
| Ventral_Stream_Visual_L | L_VMV2 | 172 | 0.27 |
| Posterior_Cingulate_L | L_31pd | 143 | 0.21 |
| Posterior_Cingulate_L | L_31a | 124 | 0.37 |
| Ventral_Stream_Visual_L | L_VVC | 182 | 0.24 |
| Anterior_Cingulate_and_Medial_Prefrontal_L | L_25 | 12 | 1.22 |
| Anterior_Cingulate_and_Medial_Prefrontal_L | L_s32 | 16 | 1.33 |
| Anterior_Cingulate_and_Medial_Prefrontal_L | L_pOFC | 21 | 0.73 |
| Insular_and_Frontal_Opercular_L | L_PoI1 | 83 | 0.42 |
| Insular_and_Frontal_Opercular_L | L_Ig | 68 | 0.46 |
| Insular_and_Frontal_Opercular_L | L_FOP5 | 84 | 0.37 |
| Orbital_and_Polar_Frontal_L | L_p10p | 131 | 0.27 |
| Inferior_Frontal_L | L_p47r | 77 | 0.54 |
| Lateral_Temporal_L | L_TGv | 111 | 0.38 |
| Early_Auditory_L | L_MBelt | 136 | 0.30 |
| Early_Auditory_L | L_LBelt | 145 | 0.29 |

| | | | |
|---|---|---|---|
| Auditory_Association_L | L_A4 | 176 | 0.24 |
| Auditory_Association_L | L_STSva | 147 | 0.30 |
| Lateral_Temporal_L | L_TE1m | 118 | 0.29 |
| Insular_and_Frontal_Opercular_L | L_PI | 82 | 0.50 |
| Anterior_Cingulate_and_Medial_Prefrontal_L | L_a32pr | 102 | 0.49 |
| Anterior_Cingulate_and_Medial_Prefrontal_L | L_p24 | 80 | 0.60 |
| Primary_Visual_R | R_V1 | 157 | 0.31 |
| MT+_Complex_and_Neighboring_Visual_Areas_R | R_MST | 182 | 0.26 |
| Dorsal_Stream_Visual_R | R_V6 | 102 | 0.37 |
| Early_Visual_R | R_V2 | 158 | 0.30 |
| Early_Visual_R | R_V3 | 169 | 0.31 |
| Early_Visual_R | R_V4 | 185 | 0.23 |
| Ventral_Stream_Visual_R | R_V8 | 184 | 0.25 |
| Somatosensory_and_Motor_R | R_4 | 87 | 0.54 |
| Somatosensory_and_Motor_R | R_3b | 90 | 0.49 |
| Premotor_R | R_FEF | 120 | 0.38 |
| Premotor_R | R_PEF | 113 | 0.40 |
| Premotor_R | R_55b | 106 | 0.40 |
| Dorsal_Stream_Visual_R | R_V3A | 135 | 0.38 |
| Posterior_Cingulate_R | R_RSC | 134 | 0.29 |
| Posterior_Cingulate_R | R_POS2 | 147 | 0.30 |
| Dorsal_Stream_Visual_R | R_V7 | 190 | 0.17 |
| Dorsal_Stream_Visual_R | R_IPS1 | 167 | 0.21 |
| Ventral_Stream_Visual_R | R_FFC | 157 | 0.34 |
| Dorsal_Stream_Visual_R | R_V3B | 168 | 0.33 |
| MT+_Complex_and_Neighboring_Visual_Areas_R | R_LO1 | 198 | 0.19 |
| MT+_Complex_and_Neighboring_Visual_Areas_R | R_LO2 | 159 | 0.35 |
| Ventral_Stream_Visual_R | R_PIT | 151 | 0.36 |
| MT+_Complex_and_Neighboring_Visual_Areas_R | R_MT | 175 | 0.25 |
| Early_Auditory_R | R_A1 | 121 | 0.32 |
| Temporo-Parieto-Occipital_Junction_R | R_PSL | 130 | 0.33 |
| Dorsolateral_Prefrontal_R | R_SFL | 127 | 0.32 |
| Posterior_Cingulate_R | R_PCV | 136 | 0.37 |
| Temporo-Parieto-Occipital_Junction_R | R_STV | 137 | 0.31 |
| Superior_Parietal_R | R_7Pm | 129 | 0.34 |
| Posterior_Cingulate_R | R_7m | 131 | 0.31 |
| Posterior_Cingulate_R | R_POS1 | 141 | 0.38 |
| Posterior_Cingulate_R | R_23d | 108 | 0.44 |
| Posterior_Cingulate_R | R_v23ab | 94 | 0.43 |
| Posterior_Cingulate_R | R_d23ab | 111 | 0.33 |
| Posterior_Cingulate_R | R_31pv | 119 | 0.28 |
| Paracentral_Lobular_and_Mid_Cingulate_R | R_5m | 57 | 0.70 |
| Paracentral_Lobular_and_Mid_Cingulate_R | R_5mv | 116 | 0.37 |
| Paracentral_Lobular_and_Mid_Cingulate_R | R_23c | 129 | 0.22 |
| Paracentral_Lobular_and_Mid_Cingulate_R | R_5L | 87 | 0.48 |
| Paracentral_Lobular_and_Mid_Cingulate_R | R_24dd | 68 | 0.52 |
| Paracentral_Lobular_and_Mid_Cingulate_R | R_24dv | 53 | 0.59 |
| Superior_Parietal_R | R_7AL | 149 | 0.32 |
| Paracentral_Lobular_and_Mid_Cingulate_R | R_SCEF | 92 | 0.41 |
| Paracentral_Lobular_and_Mid_Cingulate_R | R_6ma | 101 | 0.53 |
| Superior_Parietal_R | R_7Am | 134 | 0.38 |
| Superior_Parietal_R | R_7PL | 136 | 0.29 |
| Superior_Parietal_R | R_7PC | 157 | 0.34 |
| Superior_Parietal_R | R_LIPv | 182 | 0.20 |
| Superior_Parietal_R | R_VIP | 172 | 0.19 |
| Superior_Parietal_R | R_MIP | 139 | 0.32 |
| Somatosensory_and_Motor_R | R_1 | 75 | 0.51 |
| Somatosensory_and_Motor_R | R_2 | 119 | 0.51 |
| Somatosensory_and_Motor_R | R_3a | 83 | 0.48 |
| Premotor_R | R_6d | 69 | 0.68 |
| Paracentral_Lobular_and_Mid_Cingulate_R | R_6mp | 57 | 0.59 |
| Premotor_R | R_6v | 123 | 0.47 |
| Anterior_Cingulate_and_Medial_Prefrontal_R | R_p24pr | 78 | 0.46 |
| Anterior_Cingulate_and_Medial_Prefrontal_R | R_33pr | 58 | 0.66 |
| Anterior_Cingulate_and_Medial_Prefrontal_R | R_a24pr | 88 | 0.53 |
| Anterior_Cingulate_and_Medial_Prefrontal_R | R_p32pr | 79 | 0.62 |
| Anterior_Cingulate_and_Medial_Prefrontal_R | R_a24 | 67 | 0.52 |
| Anterior_Cingulate_and_Medial_Prefrontal_R | R_d32 | 104 | 0.49 |
| Anterior_Cingulate_and_Medial_Prefrontal_R | R_8BM | 84 | 0.58 |
| Anterior_Cingulate_and_Medial_Prefrontal_R | R_p32 | 88 | 0.52 |
| Anterior_Cingulate_and_Medial_Prefrontal_R | R_10r | 40 | 0.82 |
| Orbital_and_Polar_Frontal_R | R_47m | 35 | 0.86 |
| Dorsolateral_Prefrontal_R | R_8Av | 119 | 0.35 |
| Dorsolateral_Prefrontal_R | R_8Ad | 91 | 0.43 |
| Anterior_Cingulate_and_Medial_Prefrontal_R | R_9m | 108 | 0.35 |

| | | | |
|---|---|---|---|
| Dorsolateral_Prefrontal_R | R_8BL | 119 | 0.25 |
| Dorsolateral_Prefrontal_R | R_9p | 93 | 0.24 |
| Orbital_and_Polar_Frontal_R | R_10d | 76 | 0.45 |
| Dorsolateral_Prefrontal_R | R_8C | 85 | 0.56 |
| Inferior_Frontal_R | R_44 | 79 | 0.50 |
| Inferior_Frontal_R | R_45 | 115 | 0.29 |
| Inferior_Frontal_R | R_47l | 59 | 0.51 |
| Inferior_Frontal_R | R_a47r | 118 | 0.29 |
| Premotor_R | R_6r | 98 | 0.54 |
| Inferior_Frontal_R | R_IFJa | 68 | 0.51 |
| Inferior_Frontal_R | R_IFJp | 71 | 0.44 |
| Inferior_Frontal_R | R_IFSp | 72 | 0.40 |
| Inferior_Frontal_R | R_IFSa | 70 | 0.47 |
| Dorsolateral_Prefrontal_R | R_p9-46v | 82 | 0.54 |
| Dorsolateral_Prefrontal_R | R_46 | 124 | 0.42 |
| Dorsolateral_Prefrontal_R | R_a9-46v | 115 | 0.39 |
| Dorsolateral_Prefrontal_R | R_9-46d | 113 | 0.46 |
| Dorsolateral_Prefrontal_R | R_9a | 108 | 0.28 |
| Anterior_Cingulate_and_Medial_Prefrontal_R | R_10v | 80 | 0.64 |
| Orbital_and_Polar_Frontal_R | R_a10p | 86 | 0.49 |
| Orbital_and_Polar_Frontal_R | R_10pp | 55 | 0.54 |
| Orbital_and_Polar_Frontal_R | R_11l | 86 | 0.47 |
| Orbital_and_Polar_Frontal_R | R_13l | 35 | 0.81 |
| Orbital_and_Polar_Frontal_R | R_OFC | 11 | 1.25 |
| Orbital_and_Polar_Frontal_R | R_47s | 71 | 0.40 |
| Superior_Parietal_R | R_LIPd | 81 | 0.51 |
| Premotor_R | R_6a | 146 | 0.36 |
| Dorsolateral_Prefrontal_R | R_i6-8 | 119 | 0.43 |
| Dorsolateral_Prefrontal_R | R_s6-8 | 97 | 0.36 |
| Posterior_Opercular_R | R_43 | 109 | 0.34 |
| Posterior_Opercular_R | R_OP4 | 118 | 0.32 |
| Posterior_Opercular_R | R_OP1 | 84 | 0.47 |
| Posterior_Opercular_R | R_OP2-3 | 74 | 0.38 |
| Early_Auditory_R | R_52 | 75 | 0.42 |
| Early_Auditory_R | R_RI | 91 | 0.39 |
| Early_Auditory_R | R_PFcm | 102 | 0.44 |
| Insular_and_Frontal_Opercular_R | R_PoI2 | 93 | 0.43 |
| Auditory_Association_R | R_TA2 | 106 | 0.36 |
| Insular_and_Frontal_Opercular_R | R_FOP4 | 97 | 0.48 |
| Insular_and_Frontal_Opercular_R | R_MI | 88 | 0.49 |
| Insular_and_Frontal_Opercular_R | R_Pir | 44 | 0.65 |
| Insular_and_Frontal_Opercular_R | R_AVI | 60 | 0.75 |
| Insular_and_Frontal_Opercular_R | R_AAIC | 65 | 0.69 |
| Posterior_Opercular_R | R_FOP1 | 105 | 0.30 |
| Insular_and_Frontal_Opercular_R | R_FOP3 | 57 | 0.56 |
| Insular_and_Frontal_Opercular_R | R_FOP2 | 71 | 0.56 |
| Inferior_Parietal_R | R_PFt | 129 | 0.51 |
| Superior_Parietal_R | R_AIP | 134 | 0.42 |
| Medial_Temporal_R | R_EC | 46 | 0.59 |
| Medial_Temporal_R | R_PreS | 87 | 0.48 |
| Medial_Temporal_R | R_H | 22 | 1.05 |
| Posterior_Cingulate_R | R_ProS | 119 | 0.40 |
| Medial_Temporal_R | R_PeEc | 48 | 0.58 |
| Auditory_Association_R | R_STGa | 153 | 0.29 |
| Early_Auditory_R | R_PBelt | 147 | 0.27 |
| Auditory_Association_R | R_A5 | 184 | 0.24 |
| Medial_Temporal_R | R_PHA1 | 155 | 0.32 |
| Medial_Temporal_R | R_PHA3 | 167 | 0.27 |
| Auditory_Association_R | R_STSda | 160 | 0.24 |
| Auditory_Association_R | R_STSdp | 175 | 0.26 |
| Auditory_Association_R | R_STSvp | 158 | 0.26 |
| Lateral_Temporal_R | R_TGd | 125 | 0.29 |
| Lateral_Temporal_R | R_TE1a | 121 | 0.29 |
| Lateral_Temporal_R | R_TE1p | 78 | 0.45 |
| Lateral_Temporal_R | R_TE2a | 55 | 0.49 |
| Medial_Temporal_R | R_TF | 98 | 0.42 |
| Lateral_Temporal_R | R_TE2p | 145 | 0.30 |
| Lateral_Temporal_R | R_PHT | 90 | 0.53 |
| MT+_Complex_and_Neighboring_Visual_Areas_R | R_PH | 186 | 0.24 |
| Temporo-Parieto-Occipital_Junction_R | R_TPOJ1 | 165 | 0.20 |
| Temporo-Parieto-Occipital_Junction_R | R_TPOJ2 | 144 | 0.31 |
| Temporo-Parieto-Occipital_Junction_R | R_TPOJ3 | 118 | 0.49 |
| Posterior_Cingulate_R | R_DVT | 160 | 0.24 |
| Inferior_Parietal_R | R_PGp | 175 | 0.24 |
| Inferior_Parietal_R | R_IP2 | 106 | 0.47 |

| | | | |
|---|---|---|---|
| Inferior_Parietal_R | R_IP1 | 125 | 0.35 |
| Inferior_Parietal_R | R_IP0 | 182 | 0.28 |
| Inferior_Parietal_R | R_PFop | 138 | 0.40 |
| Inferior_Parietal_R | R_PF | 136 | 0.35 |
| Inferior_Parietal_R | R_PFm | 136 | 0.28 |
| Inferior_Parietal_R | R_PGi | 143 | 0.25 |
| Inferior_Parietal_R | R_PGs | 156 | 0.27 |
| Dorsal_Stream_Visual_R | R_V6A | 165 | 0.23 |
| Ventral_Stream_Visual_R | R_VMV1 | 124 | 0.37 |
| Ventral_Stream_Visual_R | R_VMV3 | 182 | 0.28 |
| Medial_Temporal_R | R_PHA2 | 171 | 0.27 |
| MT+_Complex_and_Neighboring_Visual_Areas_R | R_V4t | 186 | 0.28 |
| MT+_Complex_and_Neighboring_Visual_Areas_R | R_FST | 192 | 0.14 |
| MT+_Complex_and_Neighboring_Visual_Areas_R | R_V3CD | 192 | 0.27 |
| MT+_Complex_and_Neighboring_Visual_Areas_R | R_LO3 | 183 | 0.25 |
| Ventral_Stream_Visual_R | R_VMV2 | 179 | 0.30 |
| Posterior_Cingulate_R | R_31pd | 136 | 0.25 |
| Posterior_Cingulate_R | R_31a | 138 | 0.29 |
| Ventral_Stream_Visual_R | R_VVC | 185 | 0.25 |
| Anterior_Cingulate_and_Medial_Prefrontal_R | R_25 | 22 | 1.05 |
| Anterior_Cingulate_and_Medial_Prefrontal_R | R_s32 | 15 | 1.43 |
| Anterior_Cingulate_and_Medial_Prefrontal_R | R_pOFC | 12 | 1.51 |
| Insular_and_Frontal_Opercular_R | R_PoI1 | 61 | 0.51 |
| Insular_and_Frontal_Opercular_R | R_Ig | 46 | 0.59 |
| Insular_and_Frontal_Opercular_R | R_FOP5 | 62 | 0.78 |
| Orbital_and_Polar_Frontal_R | R_p10p | 118 | 0.36 |
| Inferior_Frontal_R | R_p47r | 75 | 0.39 |
| Lateral_Temporal_R | R_TGv | 85 | 0.37 |
| Early_Auditory_R | R_MBelt | 130 | 0.27 |
| Early_Auditory_R | R_LBelt | 128 | 0.28 |
| Auditory_Association_R | R_A4 | 167 | 0.26 |
| Auditory_Association_R | R_STSva | 117 | 0.39 |
| Lateral_Temporal_R | R_TE1m | 85 | 0.41 |
| Insular_and_Frontal_Opercular_R | R_PI | 62 | 0.47 |
| Anterior_Cingulate_and_Medial_Prefrontal_R | R_a32pr | 95 | 0.57 |
| Anterior_Cingulate_and_Medial_Prefrontal_R | R_p24 | 87 | 0.60 |
| Subcortical_L | L_Thalam | 63 | 0.55 |
| Subcortical_L | L_Caudate | 57 | 0.57 |
| Subcortical_L | L_Putamen | 36 | 0.85 |
| Subcortical_L | L_Pallidum | 8 | 1.29 |
| Subcortical_L | L_Hipp | 51 | 0.70 |
| Subcortical_L | L_Amygdala | 78 | 0.52 |
| Subcortical_L | L_Accumbens | 24 | 0.83 |
| Subcortical_L | R_Thalam | 48 | 0.64 |
| Subcortical_R | R_Caudate | 52 | 0.60 |
| Subcortical_R | R_Putamen | 27 | 0.95 |
| Subcortical_R | R_Pallidum | 9 | 1.36 |
| Subcortical_R | R_Hipp | 50 | 0.78 |
| Subcortical_R | R_Amygdala | 85 | 0.47 |
| Subcortical_R | R_Accumbens | 17 | 0.81 |
